# Vps4 triggers sequential subunit exchange in ESCRT-III polymers that drives membrane constriction and fission

**DOI:** 10.1101/718080

**Authors:** Anna-Katharina Pfitzner, Vincent Mercier, Aurélien Roux

## Abstract

ESCRT-III is a ubiquitous complex which catalyzes membrane fission from within membrane necks via an as yet unknown mechanism. Here, we reconstituted in vitro the ESCRT-III complex onto membranes. We show that based on variable affinities between ESCRT-III proteins and the ATPase Vps4, subunits are recruited to the membrane in a Vps4-driven sequence that starts with Snf7 and ends with Did2 and Ist1 which, together, form a fission-active subcomplex. Sequential recruitment of ESCRT-III subunits is coupled to membrane remodeling. Binding of Did2 promoted the formation of membrane protrusions which later constricted and underwent fission mediated by the recruitment of Ist1. Overall, our results provide a mechanism to explain how a sequence of ESCRT-III subunits drives membrane deformation and fission.

## Main Text

Endosomal Sorting Complex Required for Transport-III (ESCRT-III) assemblies catalyze membrane fission from within membrane necks in all investigated cellular processes requiring this type of fission event (*1–16*). ESCRT-III polymers are nucleated by several factors. ESCRT-II, which binds ESCRT-III core subunit Vps20, is the canonical one. Besides Vps20, yeast ESCRT-III complex contains three other core subunits: Snf7, which binds Vps20-ESCRT-II, as well as Vps2 and Vps24, which, in tandem, bind Snf7 and recruit the AAA-ATPase Vps4 via their MIM-domain (*1, 17–26*). While all ESCRT-III subunits have MIM-domains, their specific affinities for Vps4 differ widely (*22, 23, 25*). Vps4 ATPase activity remodels ESCRT-III by inducing turnover of ESCRT-III subunits, controlling the balance between polymer growth (*27*) and disassembly (*24–26*). ESCRT-III core subunits assemble, alone or in various stoichiometries, into single or multiple stranded filaments (*24, 28–33*). These filaments usually exhibit high spontaneous curvature, but a low rigidity, which leads to diverse helical shapes like spirals (*30, 32–34*), conical spirals (*28, 34*) and tubular helices (*24, 35*). While the possibility that transitions occur between those shapes is under debate (*36*) many findings support the notion that Vps4-dependent ESCRT-III remodeling promotes membrane constriction and fission (*7, 27, 37*). The most direct evidence is the asymmetric constriction of in vitro formed CHMP2-CHMP3 (mammalian homologs of Vps2 and Vps24) tubular copolymers by Vps4 (*38*). While attempts to reconstitute fission in vitro with ESCRT-III core subunits provided conflicting results regarding the role of Vps4 (*39, 40*), the most constricted polymers observed with Snf7, Vps2 and Vps24 exceed a radius of 10 nm (*24, 27, 30, 32, 33*), far from the theoretical limit of 3 nm required for spontaneous fission (*41*) and far from the radius of experimentally observed dynamin pre-fission intermediates (*42*). This suggests that other subunits or mechanisms are at play to reach sufficient constriction for membrane fission.

Aside from core ESCRT-III proteins, accessory subunits are thought to play a role specific to subsets of ESCRT-III functions. However, their contributions to the ESCRT-III membrane remodeling function is less understood. For example in mammalian cells, CHMP7 nucleates ESCRT-III assembly in nuclear envelope reformation (*43–45*), or CHMP1B is required for Spastin recruitment to cleave microtubules at the site of ESCRT-III-mediated abscission (*7, 46, 47*). In contrast, CHMP1 and IST1 are found in many ESCRT-dependent processes and knock-down of these proteins causes strong phenotypes in abscission (*48–51*). Likewise, knock-out of Did2 (yeast homolog of CHMP1) results in cargo sorting defects similar to those observed upon depletion of ESCRT-III core subunits (*52–54*). Hence, Did2/CHMP1 may play an important, yet unrecognized, role in ESCRT-III function.

## Results

### Vps2 and Did2 form a Snf7-binding complex

To better understand the function of Did2, we first analyzed its recruitment to ESCRT-III assemblies by incubating giant unilamellar vesicles (GUVs) or Snf7-coated GUVs with equimolar amounts of Atto565-Did2 and various ESCRT-III subunits. While Did2 alone did not bind bare membranes or Snf7-coated membrane, it was co-recruited with Vps2 to Snf7-coated GUVs (Fig. S1A-B). Addition of Vps24 enhanced Vps2-mediated Did2-recruitment to Snf7-coated GUVs but failed to recruit Did2 on its own. Consistently, Atto565-Vps2 was co-recruited with Did2 to Snf7-coated GUVs. Co-recruitment of Vps2 with Vps24 was, however, stronger than co-recruitment with Did2 and could not be further enhanced by Did2 (Fig. S1C-D). These findings suggest that Vps2 and Did2 form an Snf7-binding subcomplex, a notion supported by recent reports (*52, 55*).

To investigate if Vps2 and Did2 are able to form copolymers with Snf7, as previously shown for Vps2-Vps24 (*17, 27*), Snf7-coated liposomes were incubated with Vps2, Vps24 and Did2 and then analyzed by negative stain electron microscopy. Snf7 alone formed separated spirals mostly consisting of a single-stranded filament (Fig. 1A). As previously reported (*27*), addition of Vps2 and Vps24 to Snf7 spirals resulted in strands within filaments to come by multiple of 2, indicating a one-to-one lateral co-polymerization of Vps2-Vps24 with pre-existing Snf7 filaments (Fig. 1D-E). Addition of Vps2-Did2 caused formation of intertwined spiral-like polymers, whereas separated single-start spirals were observed when Vps2-Vps24-Did2 was added (Fig.1B-C, E; S1E-F). Interestingly, both Snf7-Vps2-Did2 and Snf7-Vps2-Vps24-Did2-polymers consisted pre-dominantly of three-stranded filaments indicating that Did2 changed the filament bundling structure.

**Fig. 1.**
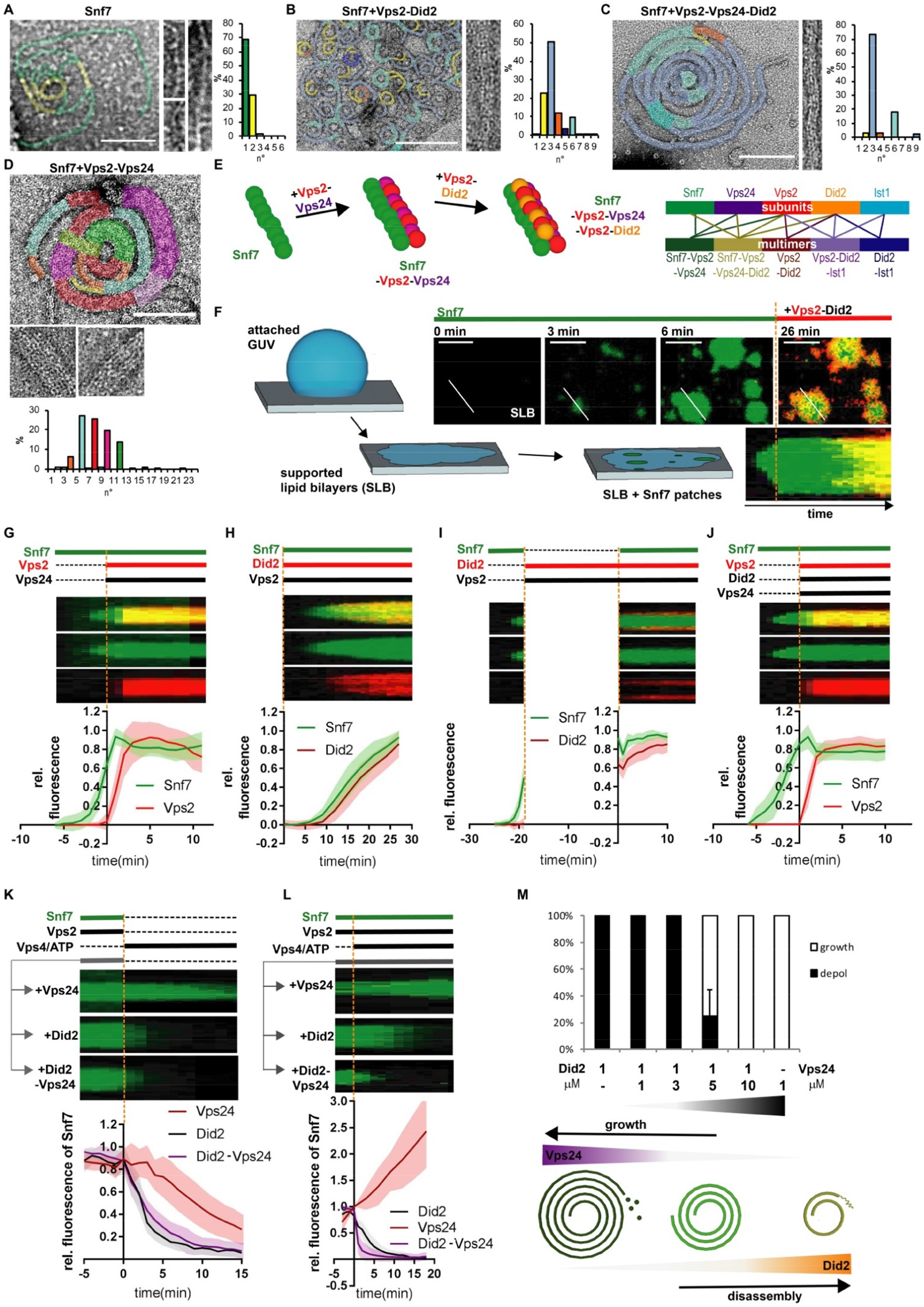
Vps2-Did2 and Vps2-Vps24 regulate Snf7 polymerization and disassembly. A-D. Negative stain electron micrographs of ESCRT-III filaments polymerized on LUVs. Histograms show the distribution of number of strands per bundle (scale bar: 100nm). E. Proposed assembly of Snf7-Vps2-Vps24-Did2 filaments. F. Scheme of supported lipid bilayer (SLB) Snf7-patch assay (scale bar: 5μm). Recombinant Alexa488-Snf7 was added at t=0 min. Recombinant Atto-565-Did2 and Vps2 were added at t=6 min. G-J. Kymographs and fluorescence quantification of Snf7-patch assays without Vps4/ATP. Color code stands for protein label, green: Alexa488, red: Atto565, black: no label. ESCRT-III proteins were added at indicated time points (orange dash line) (G: n=3 ROI=162; H: n=3 ROI=87; I: n=3 ROI=93; J: n=3 ROI=109; mean ± SD). K-L. Kymographs and fluorescence quantification patch assays with Vps4/ATP. Vps4/ATP was added at t=0 min to pre-grown Alexa488-Snf7-patches pre-incubated with indicated proteins. Alexa488-Snf7 fluorescence was measured over time (K: Vps24: n=3 ROI=94 Did2: n=4 ROI=113 Vps24-Did2: n=3 ROI=109; L: Vps24: n=3 ROI=88 Did2: n=3 ROI= 143;Vps24-Did2: n=3 ROI=142; mean ± SD). M. Percentage of growing vs depolymerizing (depol) Snf7-patches as a function of Vps24 and Did2 concentrations (n>3; mean ± SD).

Altogether our findings show that, although Snf7 is sufficient to recruit Vps2-Did2, Vps2-Vps24 enhances Did2-binding to Snf7, suggesting that Vps2-Did2 interacts with both filaments. Since we further observed that addition of Vps2-Did2 elicit three-stranded polymers, we speculated that Vps2-Vps24 initially forms a lateral copolymer along a Snf7-filament, before both strands are bridged via a Vps2-Did2 copolymer (Fig 1A-E). To test this idea of a sequential filament build-up, we next monitor the recruitment dynamics of these ESCRT-III subcomplexes using Snf7-patches grown on supported lipid bilayers (*27, 33*) (Fig. 1F).

### Vps2-Vps24 and Vps2-Did2 regulate growth and disassembly of Snf7-filaments

Snf7-patches were incubated with Vps2, Atto565-Did2 and varying concentrations of Vps24. Kymographs were extracted from midsections of patches to assess binding dynamics (Fig. 1F). Consistent with Vps24-mediated enhancement of Did2-Vps2 co-recruitment onto Snf7-coated GUVs (Fig. S1A-B), we observed that Vps2-Vps24 increased Vps2-Did2 binding rate and maximum amount, when Vps2, Vps24 and Did2 were added in a ratio of 2:1:1 (Fig. S1G-H). On the reverse, higher concentrations of Vps24 reduced binding of Atto565-Did2 to Snf7-patches. Since direct competition for binding-sites seems unlikely in a scenario where Vps24 also recruits Did2, inhibition of Did2-binding likely resulted from depletion of Vps2 from the soluble protein pool. As formation of a Did2-homopolymer on top of pre-existing Snf7-Vps2-Vps24-filaments would not be affected by depletion of free Vps2, inhibition of Did2-binding upon Vps2-depletion strongly suggests that Did2 is exclusively recruited to ESCRT-III assemblies in form of Did2-Vps2 heterodimers. We speculate that the three strands observed in Snf7-Vps2-Vps24-Did2-spirals are formed by an initial Snf7-filament along which a Vps24-Vps2-filament copolymerizes to then induces recruitment of Did2-Vps2 resulting in formation of a third Vps2-Did2-filament strand (Fig. 1C). Reversely, Did2 did not affect Vps24 binding to Snf7 (Fig. S1I).

Next, we investigated how the Vps2-Did2 complex compares to the Vps2-Vps24 complex regarding inhibition of Snf7 polymerization (*17, 27*). While addition of Vps2-Vps24 together arrested growing Snf7-patches (Fig. 1G), Snf7-patches grew slowly in presence of Vps2-Did2 (Fig. 1H). This partial inhibition of growth could be due to a lower inhibition capability of Vps2-Did2 compared to Vps2-Vps24 or resulting from a lower affinity for Snf7 polymers. To address this issue, Snf7-patches were saturated with Vps2-Did2 in absence of free Snf7, before free Snf7 was re-introduced. In this case, growth was fully inhibited (Fig. 1I). Addition of Vps2-Vps24-Did2 inhibited growth as efficiently as Vps2-Vps24 (Fig. 1J). These results indicate that the Vps2-Did2 complex, like the Vps2-Vps24 complex, has the ability to fully inhibit Snf7 polymerization. However, because of a lower affinity for Snf7 polymers, as evidenced by slower binding dynamics (Fig. S1J), Vps2-Did2, presumably, does not reach sufficient stoichiometry in the copolymer to fully inhibit Snf7 polymerization. Partial inhibition of Snf7 polymerization, likely, gives rise to the particular polymer architecture and spread distribution in number of strands per filament observed in Snf7-Vps2-Did2 polymers (Fig.1B).

The second function of Vps2-Vps24 is recruitment of Vps4, which, in turn, promotes subunit-turnover triggering either disassembly of ESCRT-III patches in absence of free Snf7 or driving their growth in presence of free Snf7 (*27*). Interestingly, Vps2-Did2 promoted disassembly of Snf7-patches more efficiently than Vps2-Vps24 (Fig. 1K) even in presence of free Snf7 (Fig. 1L). When both Vps24 and Did2 were present at equimolar ratio, Did2 dominated and promoted disassembly. Thrillingly, ESCRT-III turnover dynamics could be changed from growth to disassembly by increasing the Did2:Vps24 molar ratio in presence of free Snf7 (Fig. 1M).

These results show that Vps2-Vps24 and Vps2-Did2 subcomplexes have seemingly redundant functions in ESCRT-III assembly, with Vps2-Vps24 having a stronger affinity for Snf7-polymers, and Vps2-Did2 being more efficient in stimulating Vps4 function. Though, while being overall redundant, each subcomplex possesses functional specializations: Vps2-Vps24 blocks Snf7 polymerization, while Vps2-Did2 promotes Vps4-triggered disassembly. Since Vps2-Vps24 recruitment precedes Vps2-Did2-binding (Fig. S1J) and the Did2:Vps24 ratio controls ESCRT-III growth, we imagined a shift in ESCRT-III filament composition over time starting first with Vps2-Vps24, promoting filament assembly, followed by take-over of Vps2-Did2 which then triggers the disassembly.

### Vps4 exchanges Vps24 for Did2 within ESCRT-III polymers

To test this hypothesis, we monitored Vps4-triggered disassembly of Atto565-Vps2 or Atto565-Vps24 from Snf7-patches in presence of Did2. While Did2 accelerated Vps24-disassembly (Fig. S2A), as seen for Snf7 (Fig. 1K), it delayed Vps2-disassembly (Fig. S2B). Moreover, whereas Alexa488-Vps24 promptly disassembled upon addition of Vps4/ATP, Atto565-Did2 intensity initially increased on Snf7-patches (Fig. 2A). We, therefore, wondered whether Vps4 could exchange Vps24 with Did2 as binding partner of Vps2. Indeed, the Vps24:Vps2 ratio decreased upon Vps4-induced subunit-turnover in presence of Did2, whereas it remained stable when Did2 was missing (Fig. 2B-D). Conversely, Did2:Vps2 ratio initially increased during subunit-turnover before the ratio plateaued (Fig. 2C), implying a subunit exchange of Vps24 to Did2 (initial phase) followed by a disassembly of a then pure Did2-Vps2 filament (plateau phase). These findings suggest that, due to its stronger affinity for Snf7, the Vps24-Vps2 complex is recruited first to Snf7-patches. Vps2-Vps24, then recruits Vps2-Did2, which stimulates Vps4 activity and, thereby, removes Snf7 and Vps24, while Vps2 is stabilized at the membrane by Did2. Indeed, these notions fit well with previous *in vivo* data, describing Did2-recruitment via Vps24 and accumulation of Vps24 and Snf7 in did2Δ yeast (*53*). Overall, our data establish occurrence of specific subunit exchange of Vps24(-Vps2) with Did2(-Vps2) within persistent ESCRT-III assemblies driven by differential affinities between subunits and differing stimulatory effects on Vps4 activity. We next asked whether this subunit exchange is coupled to the membrane remodeling function of ESCRT-III.

**Fig. 2.**
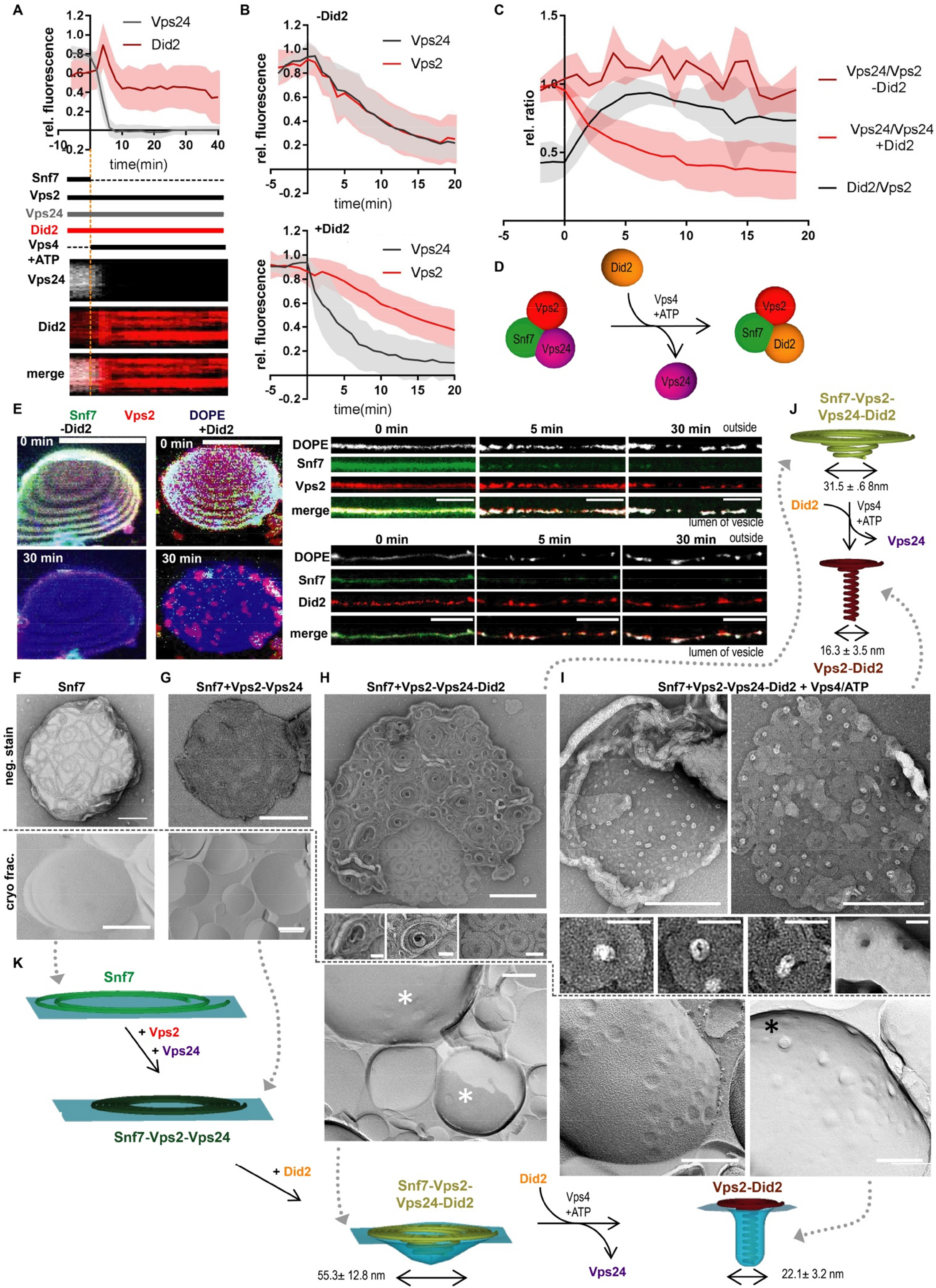
Vps4-driven exchange of Vps2-Vps24 to Vps2-Did2 triggers 3D-deformation of ESCRT-III-filaments. A. Kymographs and fluorescence quantification of patch assay in which Vps4/ATP was added at t= 0 min to pre-grown Snf7-patches pre-incubated with Alexa488-Vps24, Atto565-Did2 and Vps2. Alexa488-Vps24 and Atto565-Did2 fluorescence were plotted against time (n=3 ROI=188; mean ± SD). B. Fluorescence quantification of Snf7-patch assays in which Vps4/ATP was added at t= 0 min to pre-grown Snf7 patches pre-incubated with Alexa488-Vps24 and Atto565-Vps2 in presence or absence of Did2. Alexa488-Vps24 and Atto565-Vps2 fluorescence were plotted against time. (-Did2: n=4 ROI=173, +Did2: n=4 ROI=284; mean ± SD). C. Fluorescence ratios of Vps24/Vps2 or Did2/Vps2 plotted over time calculated from B or a respective experiment with Atto565-Did2 and Alexa488-Vps2 (Vps24/Vps2 n=4; Did2/Vps2 n=3; mean ± SD). D. Schematic representation of Vps4-triggered exchange of Vps24 to Did2. E. Tilted 3D projections of confocal Z-stacks of partially adhered GUVs (left panel; scale bar 10 μm). Vps4/ATP was added at t= 0 min to vesicles labelled with DOPE-Atto647N and pre-incubated with Alexa488-Snf7, Atto565-Vps2 and Vps24 in absence and presence of Did2. Linearized contours of GUVs (right panel; scale bar 5 μm). Vps4/ATP was added at t= 0 min to vesicles labelled with DOPE-Atto647N and pre-incubated with Alexa488-Snf7, (Atto565-)Vps2, Vps24 and (Atto565-)Did2. F-G. Negative stained micrographs of LUVs (neg. stain; upper panel) or micrographs of freeze-fractured LUVs (cryo frac.; lower panel) incubated with the indicated proteins. (scale bar 200 nm, zoom: scale bar 50 nm; white asterisk marks Did2-induced deformations; black asterisk marks fractured Did2-induced deformations). J. Schematic representation of shape changes and sizes of observed filament structures in H and I (Snf7-Vps2-Vps24-Did2: ROI=84; Snf7-Vps2-Vps24-Did2 + Vps4/ATP: ROI=111; mean ± SD). K. Schematic representation of shape changes and sizes of observed membrane structures Snf7-Vps2-Vps24-Did2: ROI=65; Snf7-Vps2-Vps24-Did2 + Vps4/ATP: ROI=54; mean ± SD).

### Did2 incorporation triggers 3D-deformation of ESCRT-III spirals

Upon Vps4-triggered disassembly of ESCRT-III polymers on GUVs in presence of Did2, we observed formation of bright puncta accumulating Atto565-Vps2, Atto565-Did2, temporarily Alexa488-Snf7 and, surprisingly, the membrane marker Atto647N-dioleoylphosphatidyl ethanolamine (DOPE) (Fig. 2E, S2C). As punctual accumulation of membranes could signify membrane deformation, liposomes coated with various ESCRT-III subunits were analyzed by negative-stain and freeze-fracture electron microscopy. Incubation with Snf7 alone, Snf7-Vps2-Vps24 or Snf7-Vps2-Vps24-Vps4/ATP yielded only flat spiral-filaments (Fig. 2F-G, S2D). Addition of Did2 resulted in moderate 3D-deformation of spiral-filaments (diameter 31.5 nm ± 6.8 nm, Fig. 2H, J) and membrane deformations (diameter 53 nm ±12.8nm; Fig. 2H, K, S2E). However, further supplementing the reaction mixture with Vps4/ATP, increasing the Did2:Vps24 molar ratio in the structures, caused massive 3D-deformations of the filaments, characterized by tubular structures protruding from the spiral center (diameter 16.3 nm ± 3.5 nm; Fig. 2I-J). Moreover, membrane deformations became more prominent and frequently extended deep inside the vesicles, eventually breaking during the fracturing procedure (Fig. 2I, K, S2E). Consistently, negative stain of the inner leaflet of burst vesicles exhibited membrane protrusions expanding towards the inside of vesicles (22.1 nm ±3.2 nm, Fig. 2I, K). These membrane protrusions are presumably formed by the observed protein filaments (16.3 nm ± 3.5 nm) and two layers of membrane bilayers (ca. 4 nm each). Altogether, we show that Did2 incorporation into ESCRT-III filaments promotes 3D-deformation of these filaments, which triggers inwards-directed membrane protrusions. Boost of membrane deformation by Vps4-driven Vps24-Did2 exchange implies that the extent of membrane deformation depends on the amounts of Did2 incorporation into the ESCRT-III structure. Furthermore, we find that Vps4 promotes rapid disassembly of Snf7 and Vps24 in presence of Did2, while Vps2-Did2 structures remain. This implies a gradual transition of subunit composition in the filament leading to a final spiral filament likely consisting predominantly of Vps2-Did2 (Fig. 1K, 2A).

### Vps2-Did2-recruited Ist1 inhibits Did2 disassembly

Did2 forms a complex with Ist1 (*56, 57*), which is also the most constricted helix of ESCRT-III filaments reported so far (*28*). We thus wondered if addition of Ist1 could trigger further constriction of the tubular protrusions observed by with Vps2-Did2. In coherence with previous *in vivo* studies (*56, 57*), incubation of Atto565-Ist1 with Snf7-patches and various ESCRT-III subunits revealed that Ist1-binding to ESCRT-III assemblies is strictly dependent on Did2-recruitment (Fig. S3A-B). Presumably, enhanced Ist1-binding observed with Vps2-Did2-Vps24 results from Vps24-mediated increase of Did2-recruitment (Fig. S1A-B). Did2-binding to Vps2 was unaffected by Ist1, excluding competitive binding (Fig. S3C).

Since Ist1 is a known modulator of Vps4 activity (*57, 58*), we tested how Ist1 impacts the disassembly of ESCRT-III polymers. Upon addition of Vps4/ATP to Snf7-Vps2-Vps24-Did2-Ist1-patches, Snf7, Vps24 and Vps2 disassembled similarly to when no Ist1 was present (Fig. 1K, 2B, 3A). In contrast, the levels of Did2 and Ist1 remained stable (Fig 3A), while Did2 and Ist1 slowly depolymerized in absence of soluble Ist1 (Fig. 3D, S3D). To further study the dynamics of this process, we switched the colors of the available soluble pools of Did2 or Ist1 during the experiments by replacing Atto488-labelled with Atto565-labelled Did2 or Ist1. We found that membrane bound Ist1 continuously exchanged with its soluble pool (Fig. 3C), in marked contrast to Did2 (Fig. 3B). Previous structural studies suggest that, due to steric hindrance, CHMP1B (mammalian homolog of Did2) can bind either IST1 or Spastin, an ATPase similar to Vps4, (*22, 23, 48, 54*). Therefore, we asked if Ist1 could prevent Did2 disassembly by masking the Vps4 binding-site of Did2. Indeed, Vps4-dependent Did2-disassembly could be abrogated by increasing the Ist1:Vps4 molar ratio (Fig. 3D, S3E), demonstrating that Ist1 and Vps4 compete for the same Did2 binding-site. Furthermore, together with Vps2 and Did2, Ist1 also accumulated in bright puncta in presence of Vps4/ATP on GUVs (Fig. S3F).

**Fig. 3.**
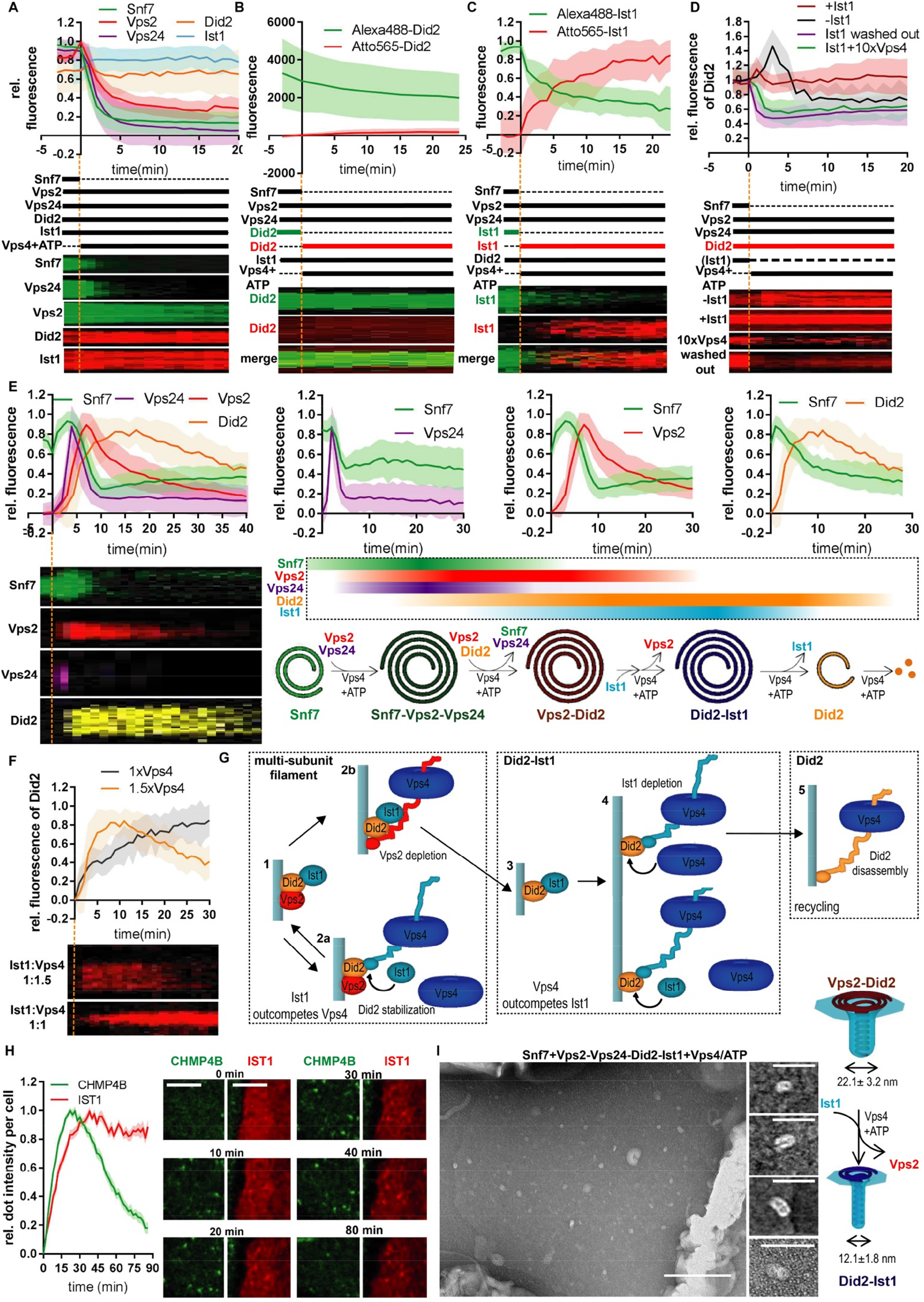
Vps4 drives a sequential recruitment and disassembly of ESCRT-III. A. Kymographs and fluorescence quantification of patch assays in which Vps4/ATP was added at t=0 min (orange dash-line) to pre-grown Snf7-patches pre-incubated with Vps24, Vps2, Did2, and Ist1. (labelled proteins: Alexa488-Snf7 n=3 ROI=108, Alexa488/Atto565-Vps2 n=3 ROI=107, Alexa488-Vps24 n=3 ROI=110, Atto565-Did2 n=4 ROI=137, Atto565-Ist1 n=4 ROI=201; mean ± SD). B. Kymographs and fluorescence quantification of patch assays in which Vps4/ATP was added at t=0 min to Snf7-patches pre-incubated with Vps24, Vps2, Alexa488-Did2, and Ist1. Before addition of Vps4, the soluble pool of Alexa488-Did2 was exchanged for Atto565-Did2. Alexa488-Did2 and Atto565-Did2 fluorescence were plotted over time (n=3 ROI=54; mean ± SD). C. Kymographs and fluorescence quantification of assays in which Vps4/ATP was added at t=0 min to Snf7-patches pre-incubated with Vps24, Vps2, Did2 and Alexa488-Ist1. Before addition of Vps4, the soluble pool of Alexa488-Ist1 was exchanged for Atto565-Ist1. Alexa488-Ist1 and Atto565-Ist1 fluorescence were plotted over time (n=3 ROI=46; mean ± SD). D. Kymographs and fluorescence quantification of patch assays in which Vps4/ATP was added at t=0 min to Snf7-patches pre-incubated with Atto565-Did2, Vps2, Vps24 and the indicated proteins. 10xVps4 correspond to a condition where Vps4 concentration was increased ten times. Atto565-Did2 fluorescence was plotted against time (-Ist1 n=3 ROI=188; +Ist1 n=4 ROI=201; 10xVps4 n= 3 ROI= 76; washed out n=3 ROI=55; mean ± SD). E. Kymographs and fluorescence quantification of patch assays and model of sequential ESCRT-III recruitment. Snf7, Vps2, Vps24, Did2, Ist1 and Vps4/ATP were added of pre-grown Alexa488-Snf7 patches at t=0 min (orange dash line). Fluorescence intensities of Alexa-488 Snf7 and Atto565-Vps2 (n=3 ROI=108; mean ± SD), Atto565-Vps24 (n=3 ROI=65; mean ± SD) and Atto565-Did2 (n=3 ROI=55; mean ± SD) were monitored and plotted against time (left panel, scaled graphs, see Fig. S3G and SI for explanations). F. Kymographs and fluorescence quantification of patch assays in which Snf7, Vps2, Vps24, Atto-565-Did2, Ist1, ATP and the indicated concentration of Vps4 were added to pre-grown Alexa488-Snf7 patches at t=0 min. Atto-565-Did2 fluorescence intensity is plotted over time (1.5xVps4: n= 3 ROI=55; 1xVps4: n=3 ROI=41; mean ± SD). G. Schematic representation of spontaneous switch from Didi2 stabilization to Did2 recycling by competition of Ist1 with other Vps4 targets. H. Fluorescence quantification and images of life-cell imaging of endosomal recruitment of CHMP4B-GFP and IST1-mcherry after hypertonic shock (HS). HS was applied at t=0 min (scale bar 5μm; CHMP4B: n=3, ROI=37; IST1: n=3, ROI=43; mean ± SEM). I. Negative stained micrographs of LUVs incubated with Snf7, Vps2, Vps24, Did2, Ist1, Vps4 and ATP and schematic representation of the observed structures (scale bar 200 nm, zoom scale bar 30 nm; (-Ist1 ROI=54; +Ist1 ROI=76; mean ± SD).

In summary, our data show that Ist1 is recruited via Did2 to the remaining Vps2-Did2 ESCRT-III assemblies, in which Ist1 specifically protects Did2 from depletion by preventing its Vps4-induced disassembly. Thus, the remaining ESCRT-III structure is essentially made of Did2 and Ist1, as all other ESCRT-III subunits, in particular Vps2, are removed by Vps4 (Fig. 3A). Our results further indicate that a cascade of subunit recruitments driven by Vps4 stimulation/inhibition, competitive inhibition and varying affinities of ESCRT-III subunits may be at play. To test this hypothesis, we set out to reconstitute this sequence of recruitment *in vitro*.

### Vps4 drives ESCRT-III machinery assembly and self-disassembly

To amplify the reaction and provide a nucleation platform for ESCRT-III assemblies, we grew Snf7-patches onto supported bilayers, to which we added a mix of Snf7-Vps24-Vps2-Did2-Ist1-Vps4/ATP in a molar ratio of 2:1:0.8:1.5:1:1.5 resembling the experimentally estimated abundance of ESCRT-III subunits (1.7:1:N/A:1.3:2.5:2.8, see (*59*) and 2.5:1:1:N/A:N/A:N/A as in (*17*)) (Fig. 3E, S3G). Immediately, Snf7 patches started to re-grow concomitant with Vps24 recruitment. During this initial phase, only modest recruitment of Vps2 and Did2 was observed. Then, Vps24 fluorescence dropped rapidly, and Snf7 patches started to disassemble, while binding of Vps2 and Did2 increased. Vps2 accumulation peaked shortly after, then Vps2 disassembled, while Did2 fluorescence still increased before dropping gradually. Importantly, Did2 disassembly proceeded only when a 1.5 Vps4:Ist1 molar ratio was added, when equal molar amounts of Ist1 and Vps4 were present, Did2 kept on saturating the polymer (Fig. 3F-G). This implies changing the balance between Ist1-dependent Did2-stabilization and Vps4-induced Did2-disassembly could allow establishment of first, a tight Did2-Ist1 polymers and then a slow disassembly of this complex. Importantly, this balance may shift during the sequential recruitment of ESCRT-III assemblies: when Vps4-targets besides Did2 and Ist1 are present, e.g. Vps2 or Vps24, Ist1-mediated Did2-protection likely outcompetes Vps4, which will deplete Vps2 and Vps24 from the polymer while Did2 is stabilized transiently at the membrane. Only once the ESCRT-III assembly, at the end of the sequence, contains solely Did2-Ist1, Vps4 can overcome Ist1-mediated protection to promote Did2-disassembly (Fig. 3G). As a support to this hypothesis, we could not detect Ist1 recruitment upon addition of the full protein mix with an Vps4:Ist1 molar ratio of 1.5, suggesting that Ist1 levels in ESCRT-III assemblies are low. However, Ist1 accumulated at ESCRT-III assemblies with an Ist1:Vps4 molar ratio of 1, mirroring incorporation of Did2 (Fig. S3H).

Our data demonstrate that ESCRT-III is a dynamic polymer that, after nucleation, automatically undergoes a Vps4-driven sequence of assembly, membrane remodeling and disassembly. We show that the initial Snf7 structure recruits primarily Vps2-Vps24 because of their high affinity for Snf7. This blocks polymerization of Snf7, but also recruits Vps4 and Did2, which dynamically exchanges with Vps24. Because Did2 further stimulates Vps4 activity, Snf7 and Vps24 rapidly disappear due to higher sensitivity for Vps4-induced depolymerization. Coinciding, Did2 accumulates, as it is protected by Ist1 from Vps4 action, while Vps2 slowly disassembles. In this sequence, we note the pivotal role of Vps2, which constitutes a backbone/bridging-element that binds all subunits (except Ist1) in the sequence.

The ESCRT-III recruitment-sequence we describe here was never observed before, probably because Did2 and Ist1, which step in later in the sequence than the core members of ESCRT-III (Vps2, Vps24 and Snf7), were omitted (*3, 37*). Upon induction of ESCRT-III recruitment to endosomes by hypertonic shock in mammalian cells (*15*), we observed a delayed recruitment of IST1 (mammalian homolog of Ist1) compared to CHMP4B (mammalian homolog of Snf7) (Fig. 3H, S4). These results evidence that the ESCRT-III recruitment sequence we found *in vitro* also occurs *in vivo* during triggered ESCRT-III activity. Following recruitment, IST1 resided at endosomes, in contrast to CHMP4B which disassembled. Since we observed similar behavior with a lowVps4:Ist1 ratio *in vitro*, we speculate that triggered global ESCRT-III recruitment might strongly diminish the cytosolic pool of Vps4 and thereby inhibit effective IST1 disassembly. *In vitro,* one cycle of ESCRT-III assembly and disassembly took several minutes, which is longer than the formation of Intra luminal vesicles (ILVs) or viral budding *in vivo* but shorter than abscission during cytokinesis (*3, 7, 60, 61*). Since the dimension of Snf7-patches and ESCRT-structures at the cytokinetic bridge largely exceed Snf7-polymers during ILV formation, we speculate that the size of ESCRT-polymer could influence the kinetics of subunit recruitment.

As a next step, we investigated how the complete sequence is coupled to the membrane remodelling functions of ESCRT-III complex, and, in particular, the impact of Ist1-binding on ESCRT-III assemblies (Snf7-Vps2-Vps24-Did2-Vps4/ATP). Strikingly, inward protrusions within liposomes measured only 12.1 nm ± 1.8 nm in presence of Ist1, suggesting that Ist1 triggers constriction of ESCRT-III polymers (Fig. 3I). Considering a thickness of 4 nm per bilayer and that membrane protrusions were measured across outer leaf flats, a distance of about 4.1 nm between inner leaf flats would be expected in these structure, which is similar to values observed in dynamin-mediated membrane fission (*42*) and is close to the theoretical estimated limit for membrane fission (*41*).

### Ist1 binding promotes membrane constriction and fission

To further characterize Ist1-mediated constriction, we took advantage of the fact that Vps2 and Did2, upon long incubation with liposomes, formed outward tubular structures with a pearled appearance. Interestingly, the diameter in the most constricted parts of these tubes (20.1 nm ± 2 nm) were comparable to the inward protrusions observed on liposomes treated with Snf7-Vps2-Vps24-Did2-Vps4/ATP (Fig. 4A, D, F compare with Fig. 2I, K). In those constricted necks, helical collars were observed (Fig. 4A, S5B).

**Fig. 4.**
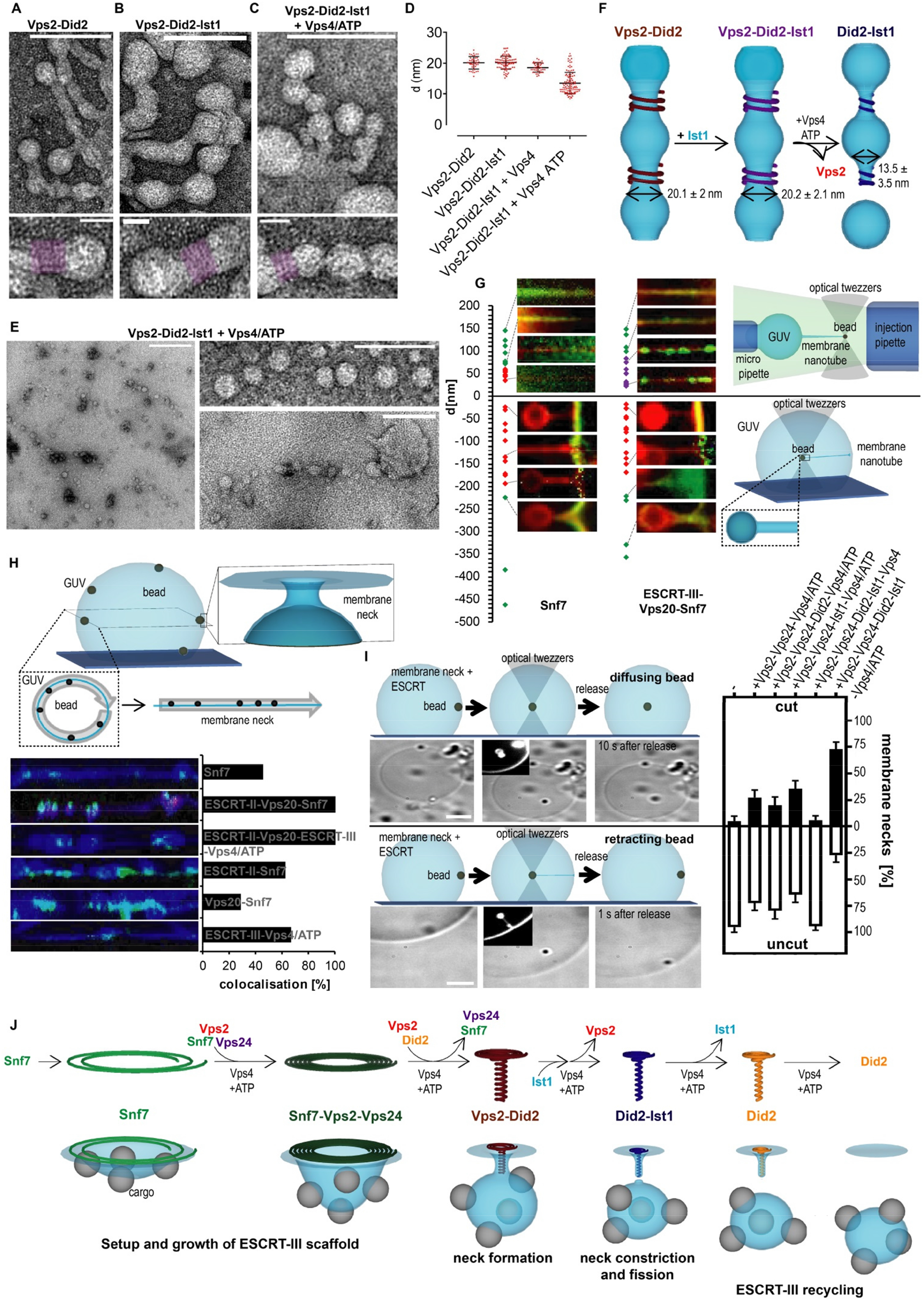
Dynamic ESCRT-III filament remodeling drives membrane fission. A-C. Negative stained micrographs of LUVs incubated with the indicated proteins (scale bar 100 nm; lower panel: scale bar 20 nm). Filament strands are indicated by overlaid purple color. D. Quantification of diameters of the helical structures (n=3; mean ± SD). E. Negative stained micrographs of LUVs pre-incubated with Vps2, Did2 and Ist1 and pre-absorbed on EM-grids before Vps4/ATP was added (scale bar 1.5 μm, zoom scale bar 100 nm). F. Schematic interpretation of the structures observed in A-E, with average radii (mean ± SD). G. Outward (upper panel) and inward (lower panel) pulled membrane nanotubes were incubated with indicated proteins. Binding of Alexa488-Snf7 was plotted against the diameter of the membrane tubes (green, binding; red, not binding; purple, nucleation of large aggregates). H. Schematic representation of artificial membrane necks assay with beads engulfed in GUVs and linearized contour of GUVs taken from confocal images of membrane necks incubated with the indicated proteins. Colocalization of Alexa488-Snf7 with artificial membrane necks (Atto647N-DOPE peaks) was quantified by measuring of the percentage of positions with both Atto647N-DOPE and Alexa488-Snf7 intensities above a threshold value along the contour (see Fig. S6C and SI). I. Schematic representation of the membrane fission assay and its discriminating criteria (see also Fig. S6H). The indicated protein mixture was added to artificial membrane necks pre-incubated with ESCRT-II, Vps20, Snf7 before necks were probed for fission by pulling beads inside the GUVs using optical tweezers and then releasing the beads (scale bar: 5μm). (-n=5 ROI=52; Snf7+Vps2+Vps24+Vps4/ATP n=4 ROI=110; Snf7 +Vps2+Vps24+Ist1+Vps4/ATP n=3 ROI=105; Snf7+Vps2+Vps24+Did2+Vps4/ATP n=3 ROI=48; Snf7+Vps2+Vps24+Did2+Ist1+Vps4 n=3 ROI=73; Snf7+Vps2+Vps24+ Did2+Ist1+Vps4/ATP n=6 ROI=133; mean ± SD). J. Model of the coupling between ESCRT-III recruitment sequence and membrane remodeling activities (see also figure S7).

Addition of Ist1 to these structures did not change the spiral diameter (20.2 nm ± 2.1 nm, Fig. 4B, D, F, S5B). However, when Vps2-Did2-Ist1 spirals were treated in solution with Vps4/ATP, polymers constricted in-between bulges (13.5 nm ± 3.5 nm; Fig.4C-D, F, S5A-B). Spiral diameters are quite spread, suggesting gradual constriction upon Vps4/ATP treatment. Based on our previous observations, Vps4 most likely depletes Vps2 from the ESCRT-polymer thereby establishing a tighter Did2-Ist1 polymer. Very excitingly, when membrane tubes were pre-absorbed onto EM grids before Vps4/ATP-treatment, constriction of Vps2-Did2-Ist1 polymers led to scission of tubules, as revealed by rows of aligned vesicles (Fig. 4E). This result may support a role for tension in ESCRT-III mediated fission, as dynamin-mediated fission could only be observed when tubes were pre-absorbed onto EM-grids using a similar assay (*62*), an effect later shown to be due to increased membrane tension (*63*).

As further evidence for ESCRT-III mediated fission, we separated liposomes and small vesicles formed during ESCRT-III action by centrifugation. When liposomes were treated with Vps2-Did2-Ist1-Vps4/ATP, a significant increase in small vesicles was observed in the supernatant compared to liposomes treated with Vps2-Did2 alone. These observations further substantiate the notion that Ist1-induced constriction leads to fission (Fig. S5C). In these experiments, however, ESCRT-III-mediated fission occurs in the reverse orientation than the native scenario, where subunits are found in the lumen of the membrane neck.

### ESCRT-II and Vps20 control orientation of ESCRT-III polymerization and fission

A powerful assay to study fission is pulling membrane tubes from GUVs (*64–66*). In this assay, streptavidin-coated beads are put in contact with an immobilized GUV containing exposed biotin moieties, and a tube is pulled by moving the bead trapped in optical tweezers. Tubes can be pulled outward the GUV, or inward the GUV (*67*). In the latter case, glass beads are nearly completely engulfed by the GUV membrane, because of the adhesive forces between glass and lipids. This assay allowed us to test recruitment of ESCRT-III assemblies as a function of positive and negative curvatures (Fig. 4G, S6A).

As reported previously, Snf7 did not polymerize on tubes pulled outwards with a radius smaller than 35 nm (Fig. 4G) (*33*). Unexpectedly, Snf7 was unable to bind to tubes pulled inward below a threshold radius of 115 nm. Addition of ESCRT-II and Vps20, which were previously suggested to trigger binding of Vps32 (*C.elegans* homolog of Snf7) or CHMP4B (human homolog of Snf7) to curved membranes (*68, 69*), partially restored binding, as large Snf7 structures formed on positively curved tubes smaller than 35 nm, but had no effect on binding of Snf7 to negatively curved tubes (Fig. 4G).

We reasoned that the unidirectional curvature of tubes might prevent Snf7 binding to negatively curved membranes. In necks, a prototypic saddle-like structure, both positive and negative curvatures are at play (Fig. 4H, S6B). To reproduce these specificities, instead of pulling engulfed glass beads into GUVs to form membrane tubes, we applied ESCRT-III subunits directly to necks connecting engulfed beads to GUV membranes (Fig. 4H). Artificial membrane neck instantly triggered, site-specific nucleation of Snf7 by ESCRT-II and Vps20, which were both sufficient and essential (Fig. 4H, S6C). Upon prolonged incubation, Snf7 alone nucleated non-specifically on GUVs.

To test if the same recruitment sequence of ESCRT-III subunits was functioning at necks as observed on supported bilayers, we added mixtures of Vps2, Vps24, Did2, Ist1 and Vps4/ATP to pre-grown ESCRT-II-Vps20-Snf7 neck-structures. Consistently, Vps2, Did2 and Ist1 were recruited to membrane necks and we observed disassembly of Snf7 and Vps2, while Did2 and Ist1 remained at membrane necks upon Vps4/ATP-induced subunit-turnover (Fig. S6D-G).

As shown above, ESCRT-III assemblies did not form onto inward-pulled membrane tubes precluding us from probing membrane fission directly. Thus, engulfed beads with pre-grown Snf7 necks were incubated with a mix of all remaining subunits (Vps2-Vps24-Did2-Ist1-Vps4/ATP), before beads were pulled inside the vesicle using optical tweezers. In this assay, we scored beads that could not be pulled into the GUV (Fig S6H), and beads that could be pulled in but when released, instantly retracted to the membrane due to a formed membrane tube, as non-fission events (Fig 4I). Only beads that, after being pulled and released, remained in the GUV center were scored as fission events (Fig.4I). In this regards it should be noted that in previous attempts to monitor ESCRT-III-mediated fission based on outward tube pulling, tube detachment due to rupture of the DOPE-biotin-streptavidin association could erroneously be confused with membrane fission–(*39*). In contrast, in our assay, internalization of beads into the GUV signified by free diffusion of beads wrapped in fluorescent membrane (Fig. 4I) could only occur by fission of the membrane neck. In absence of all ESCRT-subunits, 95.2% of the beads did not undergo fission (Fig. 4I). In contrast, fission occurred with 27.5% efficiency when the nucleators ESCRT-II and Vps20, and the core subunits of ESCRT-III Snf7, Vps2, Vps24 and Vps4/ATP, were added to GUVs. Strikingly, fission efficiency was dramatically increased to 72.8% when both Did2 and Ist1 were added to the mix (Fig. 4I). This demonstrates the importance of Did2 and Ist1 for ESCRT-III-catalyzed membrane fission. Our data strongly support the notion that sequential conversion of Snf7-Vps2-Vps24 polymers to Did2-Ist1 polymers induces enough constriction of ESCRT-III assemblies to promote efficient membrane fission.

## Discussion

Overall, our results show that Vps4-driven sequential recruitment of subunits is the mechanism by which ESCRT-III constricts membranes (Fig. 4J, S7). In clathrin-mediated endocytosis, the sequence of subunit recruitment was first identified in yeast (*70*) and later shown to be the same in mammalian cells (*71*). The ESCRT-III sequence we identified here is another example of a membrane remodeling process driven by a sequential recruitment of proteins, and had been postulated before from biochemical studies (*17, 19, 30, 40*), even though never directly observed.

We found that, in agreement with previous studies of *C.elegans* homologues (*69*), positive curvature triggers Snf7-nucleation by ESCRT-II and Vps20. In forming vesicles, positively curved membrane is limited to the rim, in contrast to negative curvature, which is found throughout the nascent bud. We, therefore, speculate that sensing positive curvature allows ESCRT-II and Vps20 to target Snf7 nucleation to the rim of shallow membrane invaginations pre-formed by crowded cargo and early ESCRT-proteins during ILV formation (*72–75*). Since Snf7 exclusively binds to low-curvature membrane and polymerizes in spiral-shape, it seems conceivable that a nucleated filament grows around the nascent bud. We envision that the initial polymer made of single-stranded Snf7 is highly flexible and with a low spontaneous curvature to adapt to large membrane necks. The sequence of recruitment probably builds stiffer assemblies through thickening and bundling (first Vps2-Vps24 and later Vps2-Did2 (see (*76*) for more detailed description)) that transform into their preferred topology and curvature which are then further constricted by addition of Ist1 and removal of Vps2.

The final Did2-Ist1 assembly generated through the described sequence is the tightest helical collar of ESCRT-III proteins characterized so far, and the only one compatible with constriction sizes that lead to fission (*28*). Besides the technical issues associated with studies in which fission was proposed to be mediated by Snf7-Vps2-Vps24 (*39, 40*), ESCRT-III structures composed of these proteins have radii of at least 14 nm (*24, 27, 30, 32–34, 77*), which is incompatible with fission. Further, we addressed the issue of ESCRT-III recycling after membrane fission. We imagine a ratio-dependent interplay of Ist1-mediated stabilization and Vps4-induced disassembly of Did2 to balance between establishing the Did2-Ist1 polymer to promote fission and recycling of the ESCRT-III structures.

Another important point solved by our study is the orientation of the protein complex towards the membrane. While in most cellular functions, the ESCRT-III complex is found inside the membrane neck it breaks, CHMP1B(Did2)-Ist1 complex can form and constrict membrane tubes from outside (*28*). This is compatible with the proposed role of ESCRT-III in fission of endosomal tubules (*78*). We observed that Did2-Ist1 can adapt and fission membrane in both orientations depending on its nucleation. Did2-Ist1 is nucleated inside membrane necks by a sequence of recruitment initiated by ESCRT-II-Vps20. If Did2-Ist1 is directly nucleated using Vps2, it forms around necks. Overall, our results identify the Did2-Ist1 complex as the most constricted ESCRT-III complex, and the one competent for fission. However, whether other important players, such as cargoes (*75*), are essential for fission remains to be investigated.

## Acknowledgments

The authors thank Nicolas Chiaruttini and Jean Gruenberg for careful correction of this manuscript and helpful discussions. AR acknowledges funding from the Swiss National Fund for Research Grants N°31003A_130520, N°31003A_149975 and N°31003A_173087, and the European Research Council Consolidator Grant N° 311536. The authors want to thank the NCCR Chemical Biology for constant support during this project.

