## Supplementary Materials for "Vps4 triggers sequential subunit exchange in ESCRT-III polymers that drives membrane constriction and fission"

**Materials and Methods**

Protein purification

ESCRT-II (Addgene plasmid #17633), Vps20 (Addgene plasmid #21490), Snf7(Addgene plasmid #21492), Vps2 (Addgene plasmid #21494), Vps24 (kind gift from James Hurley lab, UC Berkeley, USA) and Vps4 (Addgene plasmid #21495) were expressed and purified as previously described (*40*, *79*). Did2 and Ist1 (gift from David Katzmann lab, Mayo clinic, USA) were expressed and purified as previously described (*58*).

Following the labelling procedure given by the reagent provider, Snf7, Vps2, Ist1 and Vps24 were labeled with TFP-AlexaFluor-488 (Ref N°A-30005, ThermoFisher Scientific,). Vps2 and Vp24 were labeled with TFP-Atto-565 (Atto-Tec AD 565-3). Did2 and Ist1 were labeled with maleimide-Atto-565 (Atto-Tec AD 565-3). Did2 was labeled with maleimide-AlexFluor-488 (ThermoFisher Scientific, A-30005). If not otherwise mentioned, following protein concentration were used: ESCRT-II 1μM, Vps20 1μM, Snf7 500nM, Did2 1 μM, Ist1 1 μM, Vps4 1 μM, ATP 2 mM. In Figure S6E 100nM Vps2 and Vps24 were used. In experiments including Did2 Vps2 and Vps24 concentration were scaled up to 1 μM to match Did2 concentration. In general, labeled proteins were mixed 1:1 with unlabeled protein.

Preparation of giant unilamellar vesicles (GUV) and large giant unilamellar vesicles (LUV)

GUVs were prepared by electroformation: 20-30 μL of a 2mg/ml lipid solution in chloroform (DOPC:DOPS:DOPE-Atto647N:DSPE-PEG(2000)Biotin, 6:4:0.01:0.003; Avanti Polar Lipids, Atto-tec) were dried on indium-tin oxide (ITO)-coated glass slides for 1h. A growth chamber was assembled by clamping a rubber ring between the ITO-slides, filled with 500 μl of a sucrose buffer osmotically equilibrated with the experimental buffer. ITO-Slides were then connected to an AC generator set under 1V AC (10 Hz) for 1.5h. GUVs were stored at 4°C for at maximum a week.

For LUV preparation, DOPC:DOPS:Rhodamine-PE (6:4:1; 10 mg/ml) mixture was evaporated in a glass tube, 500 μl of buffer were added, the tube was vortex followed by 5 times freezing and thawing. LUVs were stored at -20 °C until usage.

Supported membrane bilayer assay and artificial membrane necks

Supported membrane bilayer assay was performed as described in (*33*). Experiments were performed in 20 mM Tris pH.6.8, 200 mM NaCl and 1mM $MgCl_2$. 2 mM DTT was added to the buffer for experiments including Ist1. GUVs diluted in buffer were burst on a plasma-cleaned coverslip forming the bottom of a flow chamber (coverslip and sticky-Slide

VI 0.4, Ibidi) to form supported bilayers. Thereafter, the chamber was passivated with Casein (1mg/ml Sigma -Aldrich) for 10 min and washed with buffer, before the experiments was conducted. Subsequent changes of protein or buffer solutions in the chamber were made via a syringe pump connected to the flow chamber.

For artificial membrane necks, partially adhered vesicles were prepared as described in (*33*). Briefly, a flow chamber assembled from a coverslip and sticky-Slide VI 0.4, Ibidi was incubated with Avidin (0.1 mg/ml) for 10 min, before washing with buffer (20 mM Tris pH.6.8, 200 mM NaCl and 1mM $MgCl_2$) and addition of GUVs (including 0.03 % DSPE-PEG(2000)Biotin) diluted in buffer. As soon as GUVs started to attach biotinylated-Albumin (1mg/ml, Sigma-Aldrich) was added to stop attachment and prevent bursting of the GUVs. After 15 min of incubation, the chamber was washed with 3 chamber volumes of buffer and glass beads diluted in buffer were added (1µm beads, 1:500; 2µm beads, 1:200; Bangs Laboratories).

For artificial necks preparation for EM, LUVs, formed in 500 mOsm sucrose, were diluted 1:100 in buffer (20 mM Tris pH.6.8, 200 mM NaCl and 1mM $MgCl_2$), spun down (10’, 5,000g) and resuspended in buffer containing 27 nm silicated $NH_2$-PS beads (1:10). Mixture was then vortexed for 30 s and incubated for 1h at RT before loading on a 75% Sucrose cushion in a centrifuge tube, topped with buffer and centrifuged 20’, 5,000g to separate bead-containing vesicles from excessive beads. The floating layer containing the Rhodamine-labeled vesicles was carefully harvested, washed, and resuspended in buffer containing 30% glycerol and processed identical to freeze-fracture samples.

Silication of Amine-polystyrene beads

40.5 ml isopropanol and 8 ml dd$H_2O$ were mixed and pH was adjusted to 11.3 using ammonia solution. 27 nm $NH_2$-PS beads (Bangs Laboratories) were sonicated (5 min), before 200 µl beads and 193 µl Tetraethylorthosilicat (TEOS) were added to the isopropanol/water mixture under stir. After incubation for 1h, beads were spun down 15’ 13,000g, washed twice with water and finally resuspended in 0.5 ml buffer. Beads were stored at 4 °C until usage.

Imaging and data analysis

Confocal Imaging was performed on an inverted spinning disc microscope assembled by 3i (Intelligent Imaging Innovation) consisting of a Nikon base (Eclipse C1, Nikon), a 100x 1.49 NA oil immersion objective and an EVOLVE EM-CCD camera (Ropper Scientific Inc.). For analysis of supported bilayer experiments, 3 µm thick Z-stack were maximally projected using a Fiji plugin (*80*). X-y drift of the microscopy was corrected using the plugin Turboreg and a custom-written ImageJ macro. For quantification, integrated fluorescence intensity of single patches was measured, normalized to either their maximum or time point 0 and a kymograph was extracted. For artificial membrane neck experiments, 15 µm thick Z-stacks were acquired. Sections containing membrane necks were selected manually, linearized and integrated fluorescence intensity was measured along the contour and/or through time using Fiji (Fig. 1C). Fluorescence intensities were normalized by their maximum value. To determine the colocalization of Snf7-Alexa488 and DOPE-Atto647N peaks, relative fluorescence was measured along linearized membrane contours after smooth averaging, relative fluorescence values were binarized (1 above threshold, 0 below, thresholds: 0.3 Snf7, 0.4 DOPE) The percentage of no colocalization was extracted by the proportion of pixels with the value 1 from the Snf7 channel for which the value in the membrane channel was 0. No colocalization was only counted at a minimal distance of four pixels to the nearest membrane neck (value 1).

Electron microscopy

For EM experiments involving Snf7, LUVs were diluted 1:100 in buffer (20 mM Tris pH.6.8, 200 mM NaCl and 1mM $MgCl_2$), spun down (10', 5,000g), resuspended in 4.5µM Snf7 for 6h (4°C), before 1 µM Vps2, 1 µM Vps24, 2 µM Did2 and 2 µM Ist1 were added overnight at 4°C. In experiments with Vps4, 1 µM Vps4, 2 mM ATP and 1 mM $MgCl_2$ were added to the samples the next morning, and incubated at 30°C for 30 min. To stop Vps4 activity, samples were chilled on ice and diluted 1:100 in buffer containing 50 mM EDTA. Finally, all samples were spun down 10' 5,000g and resuspended in buffer (negative stain EM) or buffer containing 30% glycerol (cryofreeze-fracture). Negative stain samples were absorbed onto EM grids and stained with 2% uranyl acetate for 30s. Freeze fracture samples were transferred onto sample stamps, flash-frozen and processed using a 060 Freeze-Fracture System (BAF). To grow Vps2-Did2(-Ist1) membrane tubes, 50 µl LUVs (10mg/ml) were diluted in buffer, spun 10' 5,000g and incubated with 6 µM of Vps2, Did2 and Ist1 for 60h (4°C). Samples were treated with Vps4/ATP as follow: 1 µM Vps4, 2 mM ATP and 1 mM $MgCl_2$ were either added in solution for 30 min at 30°C to 1:5 diluted samples or to 1:20 diluted samples absorbed for 5 min onto an EM-grid. Depolymerization in solution was stopped as described above. Depolymerization on grids was stopped by blotting and staining. Images were acquired on a Tecnai G2 Sphera (FEI) electron microscope.

Optical tweezer tube pulling experiment

Membrane nanotube pulling experiments were performed on the setup published in (*33*) allowing simultaneous optical tweezer application, spinning disc confocal and brightfield imaging based on an inverted Nikon eclipse Ti microscope and a 5W 1064nm laser focused through a 100 x 1.3 NA oil objective (ML5-CW-P-TKS-OTS, Manlight). Outward directed membrane nanotubes were pulled with streptavidin beads (3.05 µm, Spherotec) from a GUV containing 0.01% DSPE-PEG(2000)Biotin and aspired in a motorized micropipette (MP-285, Sutter Instrument). Proteins were injected using a slightly bigger micropipette connected to a pressure control system (MFCS-VAC -69 mbar, Fluigent). 2 µm glass beads internalized into GUVs adhered on the bottom of flow chambers (see artificial membrane necks protocol) were pulled inward to form membrane nanotubes by moving the stage relative to the bead trapped in optical tweezers. Proteins were added via a syringe pump connected to the flow chamber. Radius r of outward pulled membrane nanotubes were calculated from the force F and membrane rigidity $\kappa$ ($\kappa$ =12 kT) using the formula $r = (2\pi\kappa)/F$. F was determined following Hook's law $F= k*\Delta x$ using the bead displacement and the trap stiffness k (k=79 pN/nm/W). Radii of inward pulled tubes were estimated using mean fluorescence intensity of the tubes and a calibration curve (Fig S2A) which was established from outward pulled tubes. By changing the aspiration pressure in the pipette, the tension, and the radius of the tube can be changed.

<u>In vitro reconstitution of ESCRT-III sequence</u>

To reconstitute the ESCRT-III sequence, Snf7-patches were pre-grown on membrane bilayers and washed with 3 chamber volumes of buffer. Then, 2 µM Snf7-Alexa 488, 1 µM Vps24, 800 nM Vps2, 1.5 µM Did2, 1 µM Ist1, 1.5 µM Vps4 and 10 mM ATP were added to the pre-grown patches. To ensure maximal activity of the ATPase throughout the reconstitution, we used a higher ATP (10 mM) concentration than typically found in cells (2 mM ATP). We did not observe noticeable changes in the kinetics of any protein as compared to experiments done at 2mM, besides a faster Did2 disassembly in conditions where it disassembled. Vps2-Atto565, Vps24-Atto565 or Did-Atto565 were mixed 1:1 with unlabeled protein. Vps2, Vps24 and Did2 dynamics were imaged separately, but together with Snf7-Alexa 488 to provide a time reference. To align the experiments on the time axis a scaling factor was calculated from the fluorescence intensity curve of Snf7: all experiments were rescaled with this factor proportional to the time between the peak of Snf7 intensity, and the time at which it reached the lower plateau value after depolymerization and the distance between these time points (Fig. S4G). Vps24, Vps2 and Did2 graphs were aligned using the calculated scaling factor (Fig. 3E).

Biochemical membrane fission assay

Membrane fission assay was adapted from (*81*). Briefly, Vps2-Did2(-Ist1) membrane tubes were grown as described above. After incubation with Vps4 and ATP, liposomes were spun 15' at 250.000g, 4°C, supernatant and pellet were mixed with sample buffer and separated using SDS-PAGE at 4°C. Gels were stained with 0.1% Coomassie in 10% acetic acid and destained in dd$H_2O$. Bands were quantified using ImageJ.

Membrane neck fission assay

Artificial membrane necks generated by 1µm beads were incubated first with ESCRT-II, Vps20 and Snf7 for 5', followed by 15' incubation with Vps2, Vps24, Did2, Ist1, Vps4 and ATP, before the protein-mixture was replaced by buffer. Beads were pulled towards the center of the GUV with an optical trap and then released. Beads connected to the membrane of the GUV instantly retracted after release, in contrast to unconnected beads which stay in the center of the GUV.

Kinetics of CHMP4B and IST1 endosomal relocalization after hypertonic shock

For live-cell imaging, HeLa Kyoto cells stably expressing LAT-CHMP4B-GFP or LAT-IST1-mcherry were seeded at 1 to 1 ratio into 35mm MatTek glass bottom microwell dishes (MatTek Corporation). Before imaging, cells were rinsed 2-fold with 1ml Leibovitz's live imaging medium (Life Technologies, Thermofisher) so that cells could be incubated without CO2 equilibration at room temperature. Cells were imaged using a 100x 1.4 NA oil DIC Plan-Apochromat VC objective (Nikon) with a Nikon A1 scanning confocal microscope at speed of 1 frame every 2 minutes. Hypertonic shock was done by addition of 0.5M sucrose containing Leibovitz's medium for a final osmolarity of 900 mOsm. For the analysis, the spots were detected for each frame for both channels using the spot detector plugin in Icy software and exported as ROIs. For each timepoint the sum of dots mean intensity was calculated using Excel, the value obtained before the shock was subtracted and the curve was normalized to 1 (for its maximum value). The graph and statistics were done using Prism 8 (GraphPad software).

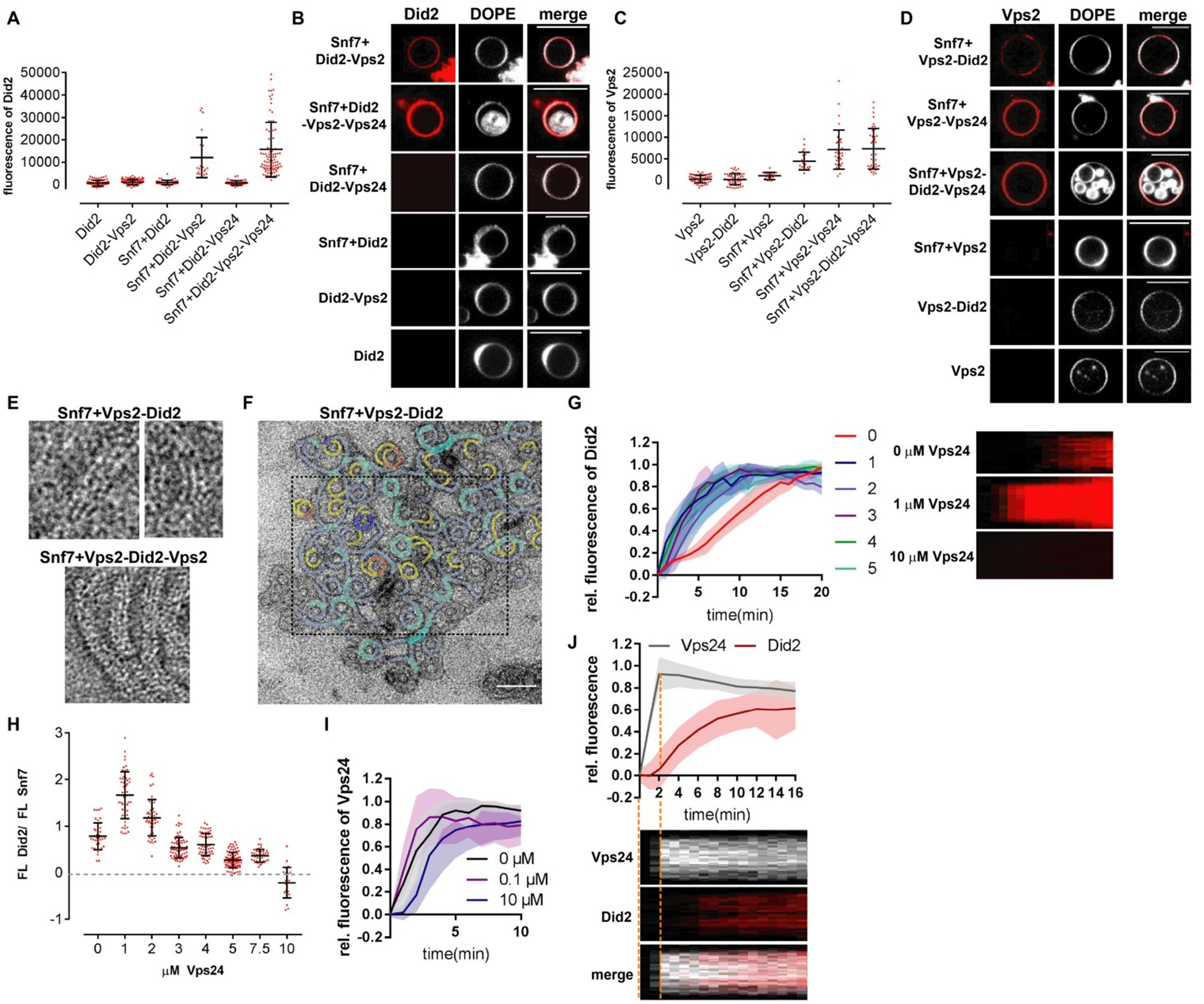


**Fig. S1.Vps2 and Did2 form a Snf7-binding complex which is recruited by Vps2-Vps24.** A-B. Confocal images and quantification of Atto565-Did2 fluorescence of LUVs or Snf7-covered LUVs incubated with Atto565-Did2 and the indicated proteins (scale bar 10μm n≥3; mean ± SD). C-D. Confocal images and quantification of Atto565-Vps2 fluorescence of LUVs or Snf7-covered LUVs incubated with Atto565-Did2 and the indicated proteins (scale bar 10μm n≥3; mean ± SD). E. Negative stain electron micrographs of ESCRT-III filaments polymerized on LUVs. Zoom of filaments described in Fig. 1B and C. F. Negative stain electron micrographs of Snf7-Vps2-Did2 filaments polymerized on LUVs. Framed area is shown in Fig. 1B (scale bar: 100 nm). G. Kymographs and fluorescence quantification of patch assay in which Atto565-Did2, Vps2 and indicated amount of Vps24 were added at t=0 min to pre-grown Alexa488-Snf7 patches. Atto565-Did2 fluorescence intensity was plotted against time (n=3, ROI≥55; mean ± SD). H. Ratio of Atto565-Did2 to Alexa488-Snf7 fluorescence intensities at plateau level from patches described in G was plotted against Vps24 concentration. I. Fluorescence quantification of Snf7 patch assays in which Alexa488-Vps24, Vps2 and indicated amount of Did2 were added at t=0 min to pre-grown Snf7 patches. Alexa488-Vps24 fluorescence intensity was plotted against time (n=3; mean ± SD). J. Kymographs and fluorescence quantification of patch assay in which Alexa488-Vps24,

Atto565-Did2 and were added at t= 0 min to pre-grown Snf7-patches Alexa488-Vps24 and Atto565-Did2 fluorescence intensities were plotted against time (n=3 ROI=188; mean ± SD).

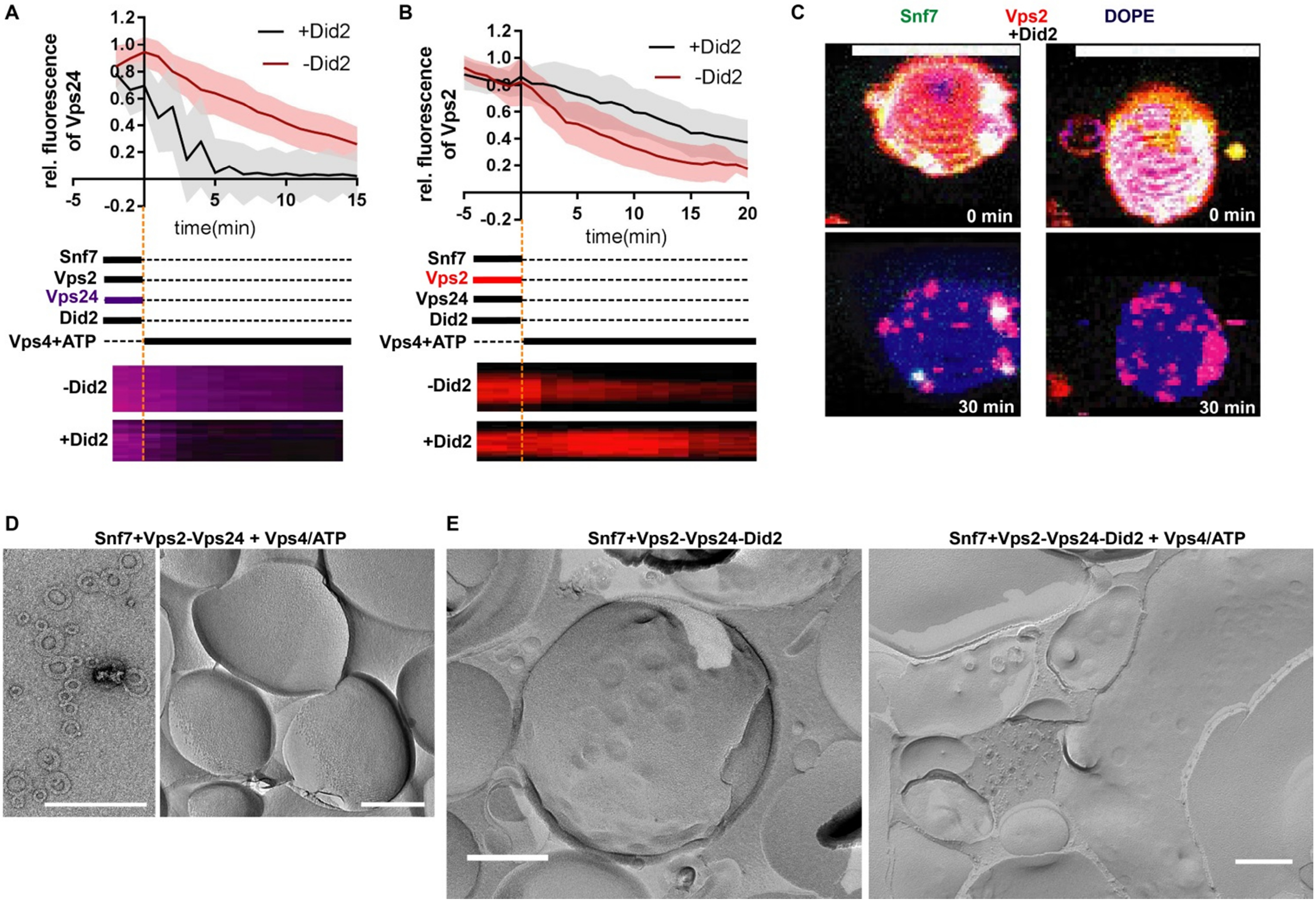


**Fig. S2. Exchange of Vps2-Vps24 to Vps2-Did2 deforms ESCRT-III-filaments.** A. Kymographs and fluorescence quantification of patch assay in which Vps4/ATP was added at t= 0 min to pre-grown Snf7-patches pre-incubated with Alexa 488-Vps24 and Vps2 in presence or absence of Did2. Alexa 488-Vps24 fluorescence intensity was plotted against time (-Did2: n=4, ROI=159; +Did2: n=3, ROI=95; mean ± SD). B. Kymographs and fluorescence quantification of patch assay in which Vps4/ATP was added at t= 0 min to pre-grown Snf7 patches pre-incubated with Vps24 and Atto565-Vps2 in presence or absence of Did2. Atto565-Vps2 fluorescence intensity was plotted against time. (-Did2: n=4, ROI=102; +Did2: n=3, ROI=137; mean ± SD). C. Tilted 3D projections of confocal Z-stacks of partially adhered GUVs (left panel; scale bar 10 µm). Vps4/ATP was added at t= 0 min to vesicles labelled with DOPE-Atto647N and pre-incubated with Alexa488-Snf7, Atto565-Vps2, Did2 and Vps24. D. Negative stained micrographs of LUVs (left panel) or micrographs of freeze-fractured LUVs (right panel) incubated with the indicated proteins. (scale bar 200 nm). E. Micrographs of freeze-fractured LUVs incubated with the indicated proteins. (scale bar 200 nm).

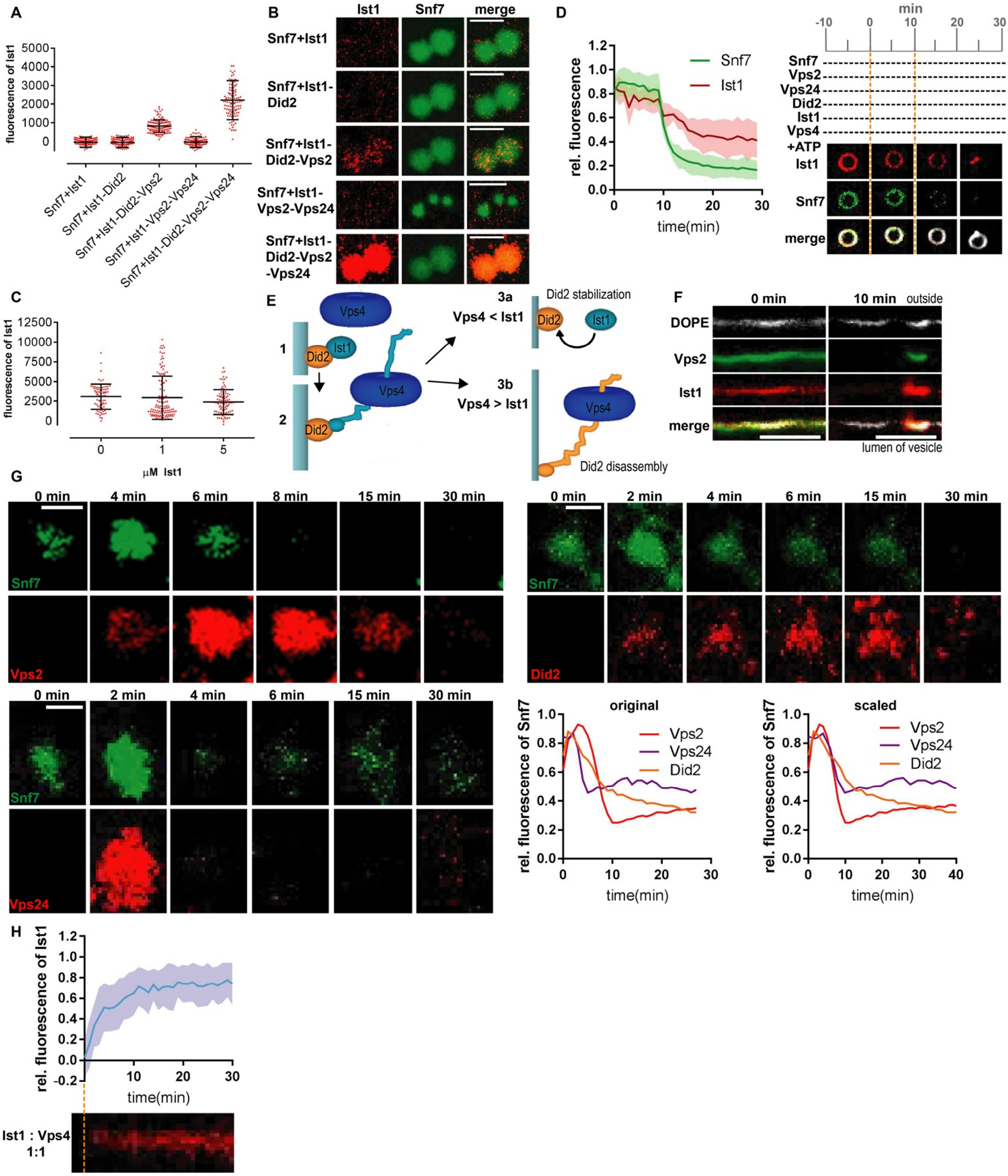


**Fig. S3. Ist1 recruitment and function in ESCRT-III assemblies.** A-B. Confocal images and quantification of Atto565-Ist1 fluorescence (n=3, mean ± SD) of patch assay in which pre-grown Alexa488-Snf7 patches were incubated with Atto565-Ist1 and the indicated proteins (scale bar 5 μm). C. Quantification of Atto565-Did2 fluorescence of patch assays in which pre-grown Snf7 patches were incubated with Atto565-Did2 and the indicated concentration of Ist1 (n=3; mean ± SD). D. Confocal images and fluorescence quantification of Snf7-coated GUVs pre-incubated with Atto565-Ist1, Vps2, Vps24 and Did2 to which buffer was added at t= 0 min and Vps4/ATP at t= 10 min (n=3 ROI=25; mean ± SD). E. Schematic representation of competition of Vps4-induced Did2-disassembly and Ist1-mediated Did2-stabilization. F. Linearized contours of GUVs. Vps4/ATP was added at t=0 min to GUVs pre-incubated with Alexa488-Vps2, Snf7, Atto565-Ist1, Did2, Vps2 andVps24. G. Confocal images of experiments described in Fig. 3E. Snf7, Vps2, Vps24, Did2, Ist1, Vps4 and ATP were added to pre-grown Alexa488-Snf7 patches at t=0 min. Fluorescence of Alexa488-Snf7 and Atto565-Vps2 (n=3 ROI=108; mean ± SD; scale bar 2 μM), or Alexa488-Snf7 and Atto565-Vps24 (n=3 ROI=65; mean ± SD; scale bar 2 μM), or Alexa488-Snf7 and Atto565-Did2 (n=3 ROI=55; mean ± SD; scale bar 2 μM) were monitored and normalized fluorescence intensities of Alexa488-Snf7 and Atto565-Vps2, Alexa488-Snf7 and Atto565-Vps24, and Alexa488-Snf7 and Atto565-Did2 were plotted against time (see Fig. 3E). Alexa488-Snf7 fluorescence graphs corresponding to the measurements with Atto565-Vps2, Atto565-Vps24 and Atto565-Did2 were aligned and scaled along the time axis (see methods for further description). Obtained scaling factors were applied on the respective Vps2, Vps24 and Did2 fluorescence graphs and scaled normalized fluorescence intensities of Snf7, Vps2, Vps24 and Did2 were plotted against time (see Fig. 3E). H. Kymographs and fluorescence quantification of patch assays in which Snf7, Vps2, Vps24, Did2, Atto-565-Ist1 and Vps4/ATP (Vps4:Ist1 ratio= 1) was added at t= 0 min to pre-grown Snf7 patches Atto-565-Ist1 fluorescence intensity was plotted over time (n=3 ROI=47; mean ± SD).

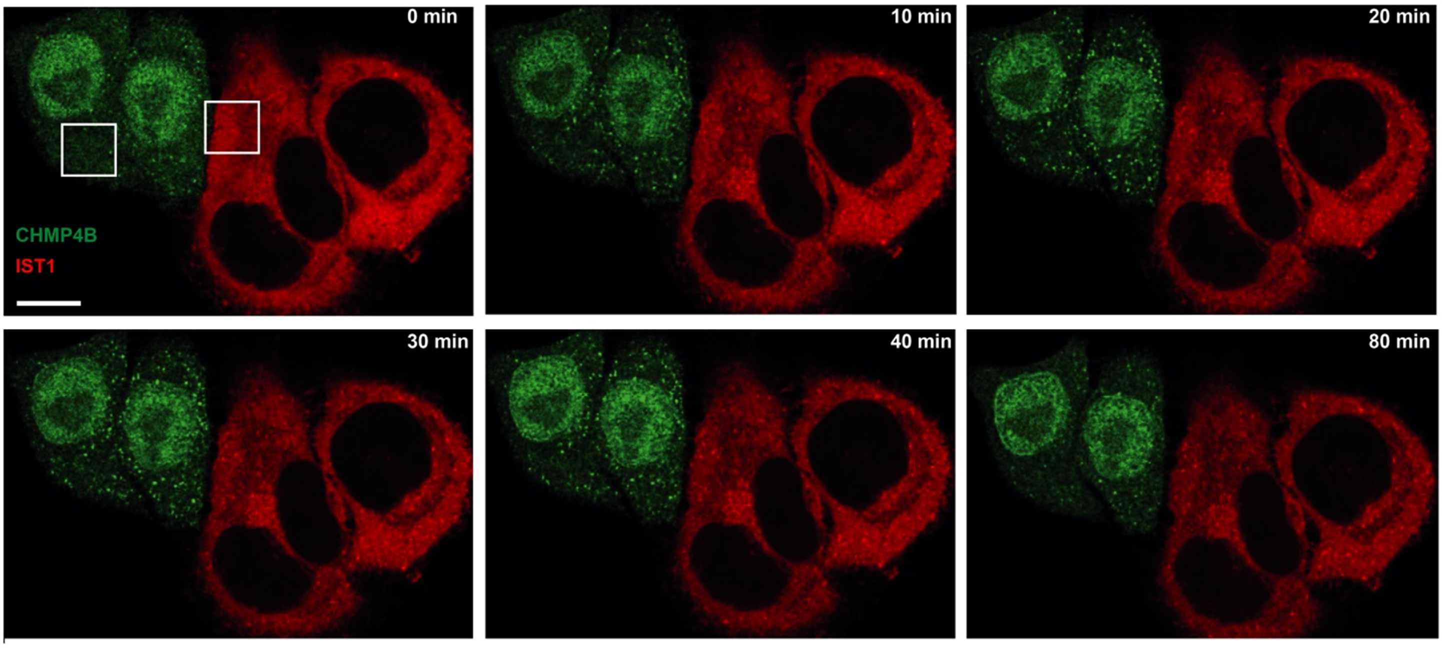


**Fig. S4. In vivo recruitment of CHMP4B and IST1.** Confocal images of live-cell imaging of endosomal recruitment of CHMP4B-GFP and IST1-mcherry after hypertonic shock (HS). HS was applied at t=0 min. ROI shown in Fig. 3H are indicated by the white boxes (scale bar 10µm; CHMP4B: n=3, ROI=37; IST1: n=3, ROI=43).

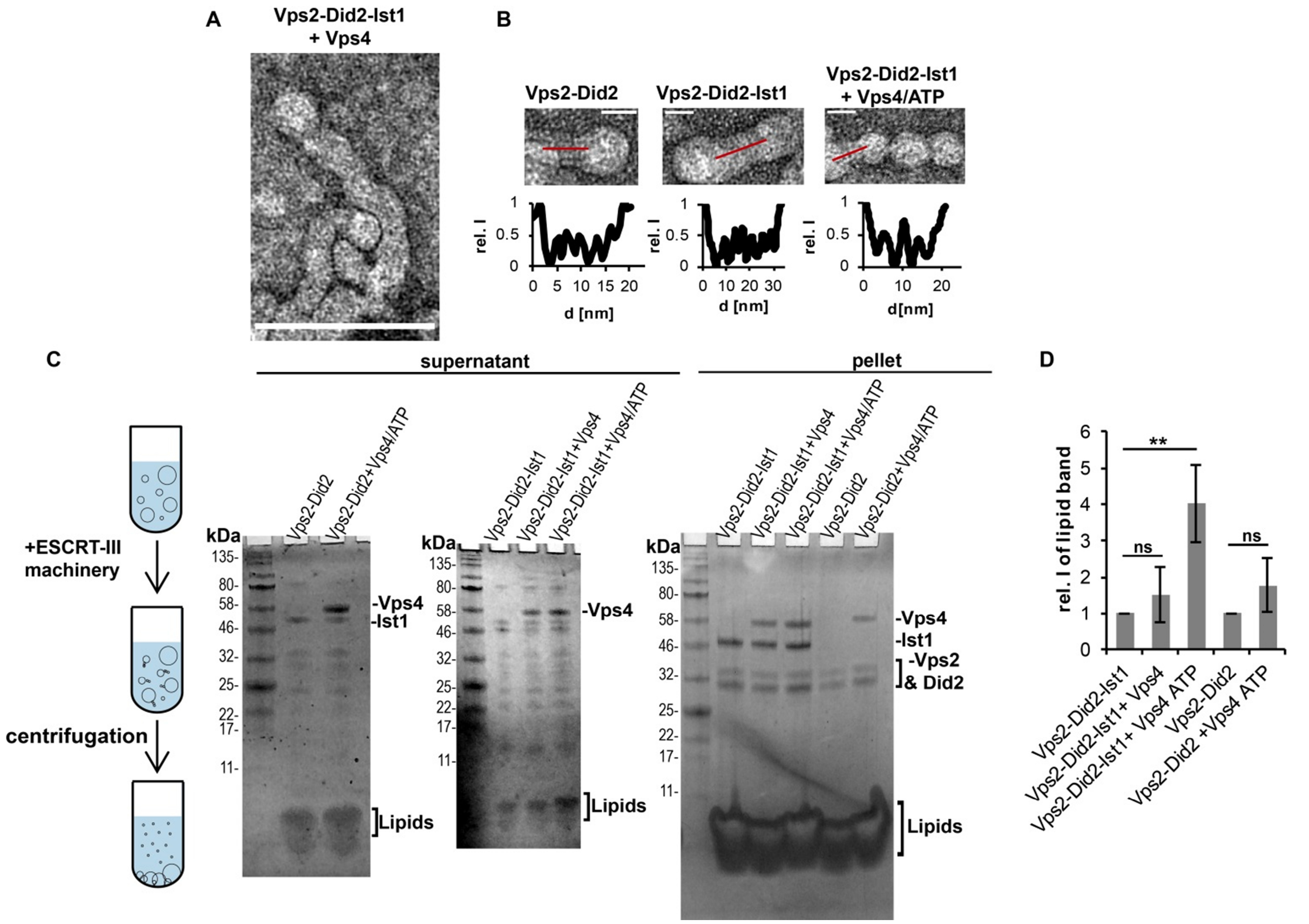


**Fig. S5. Did2-Ist1 polymers drives ESCRT-filament constriction and membrane fission.** A. Negative stained micrographs of LUVs incubated with the indicated proteins (scale bar 100 nm) B. Negative stained micrographs of LUVs incubated with the indicated proteins described in Fig.4 A-C (scale bar 20 nm). Filament profiles were analyzed by measuring intensities along the overlaid line. C. Coomassie-staining of SDS-PAGE loaded with the supernatant and pellet after ultra-centrifugation of LUVs incubated with the indicated proteins. D. Quantification of lipid-bands intensity of experiments described in C (n=4; mean ± SD, ** p-value<0.01).

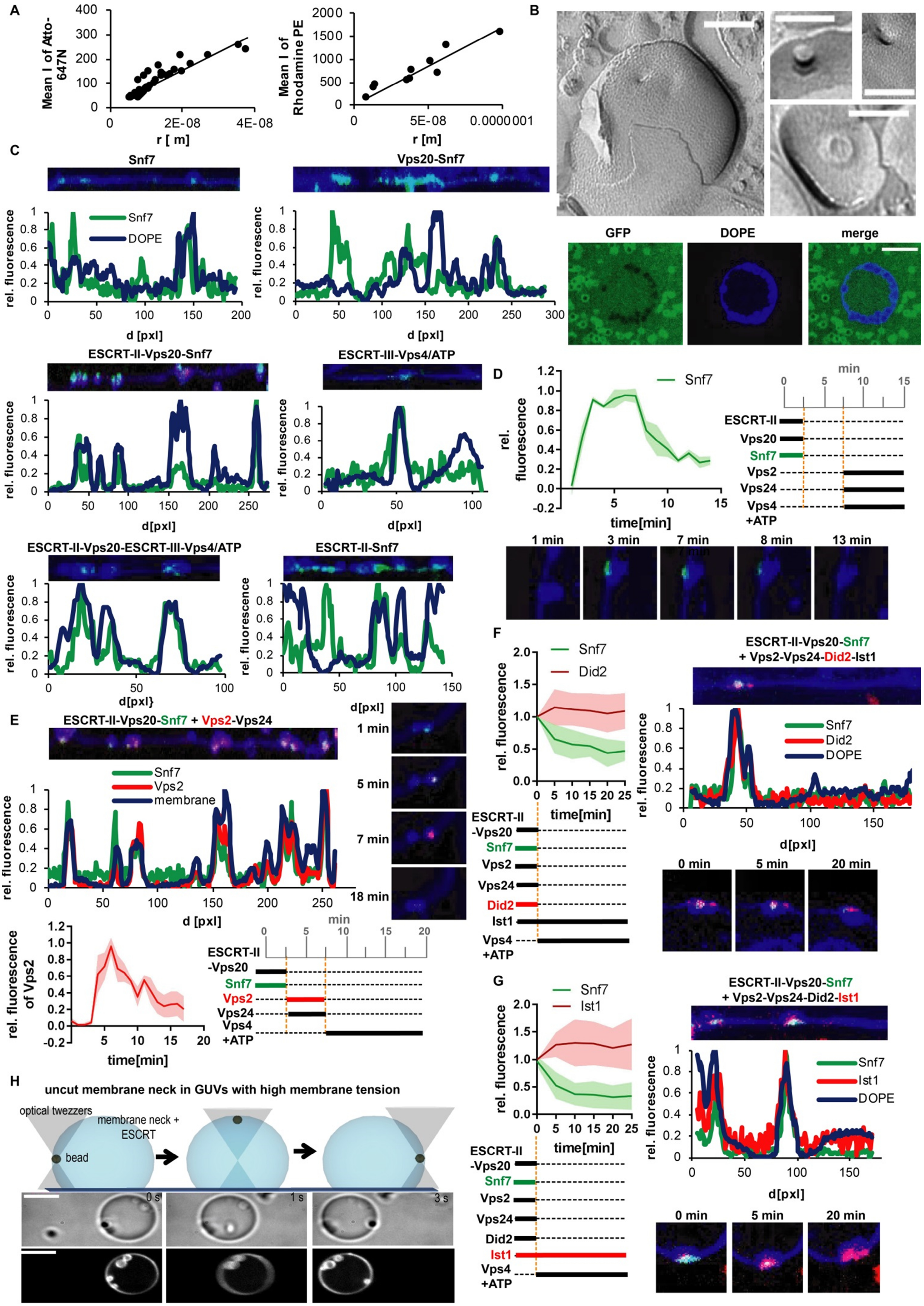
A
Mean I of Atto-647N
r [ m]
Mean I of Rhodamine PE
r [m]
B
GFP
DOPE
merge
C
Snf7
Vps20-Snf7
DOPE
rel. fluorescence
d [pxl]
ESCRT-II-Vps20-Snf7
ESCRT-III-Vps4/ATP
ESCRT-II-Vps20-ESCRT-III-Vps4/ATP
ESCRT-II-Snf7
D
rel. fluorescence
time[min]
min
ESCRT-II
Vps20
Snf7
Vps2
Vps24
Vps4
+ATP
1 min
3 min
7 min
8 min
13 min
E
ESCRT-II-Vps20-Snf7 + Vps2-Vps24
Vps2
membrane
5 min
18 min
rel. fluorescence of Vps2
ESCRT-II
-Vps20
F
Did2
ESCRT-II-Vps20-Snf7
+ Vps2-Vps24-Did2-Ist1
0 min
20 min
Ist1
G
H
uncut membrane neck in GUVs with high membrane tension
optical twezzers
membrane neck +
ESCRT
bead
0 s
1 s
3 s

**Fig. S6. ESCRT-II and Vps20 site-specifically nucleates ESCRT-III polymers at artificial membrane necks.** A. Calibration curves of lipid fluorescence mean intensity along outward-directed membrane nanotubes plotted against radii calculated fromoptical trap force value. B. Micrographs of freeze-fractured LUVs containing artificial membrane necks (upper panel; scale bar 100 nm) and confocal images of GUV containing artificial membrane necks incubated with GFP. (lower panel; scale bar 5µm) C. Linearized contour of GUVs made from confocal images of membrane necks incubated with the indicated proteins. Colocalization were analyzed by measuring the normalized intensities of Atto647N-DOPE and Alexa488-Snf7 along the contours (see Fig. 4H and methods). D. Confocal images and fluorescence quantification of artificial membrane necks incubated with the indicated sequence of proteins. Alexa488-Snf7 fluorescence intensity was measured over time (n=3, ROI=5; mean ± SD). E. Confocal images and fluorescence quantification of artificial membrane necks incubated with the indicated sequence of proteins. Atto565-Vps2 fluorescence intensity was measured over time (n=3, ROI=6; mean ± SD). F. Linearized contour of GUVs, confocal images and fluorescence quantification of artificial membrane necks incubated with the indicated sequence of proteins. Vps4/ATP was added at t=0 min to pre-grown Alexa488-Snf7 structures pre-incubated with Atto565-Did2, Vps2, Vp24 and Ist1. Normalized intensities of Atto647N-DOPE, Alexa488-Snf7 and Atto565-Did2 was measured along the contours and over time (n=3, ROI=20; mean ± SD). G. Linearized contour of GUVs, confocal images and fluorescence quantification of artificial membrane necks incubated with the indicated sequence of proteins. Vps4/ATP was added at t=0 min to pre-grown Alexa488-Snf7 necks structures pre-incubated with Atto565-Ist1, Vps2, Vp24 and Did2. Normalized intensities of Atto647N-DOPE, Alexa488-Snf7 and Atto565-Ist1 was measured along the contours and over time (n=3, ROI=23; mean ± SD). H. Schematic representation of the case of no fission event at high membrane tension. The bead can still be moved along the GUV surface, but not pulled inward. Experiments were performed as described in Fig. 4I (scale bar: 5µm).

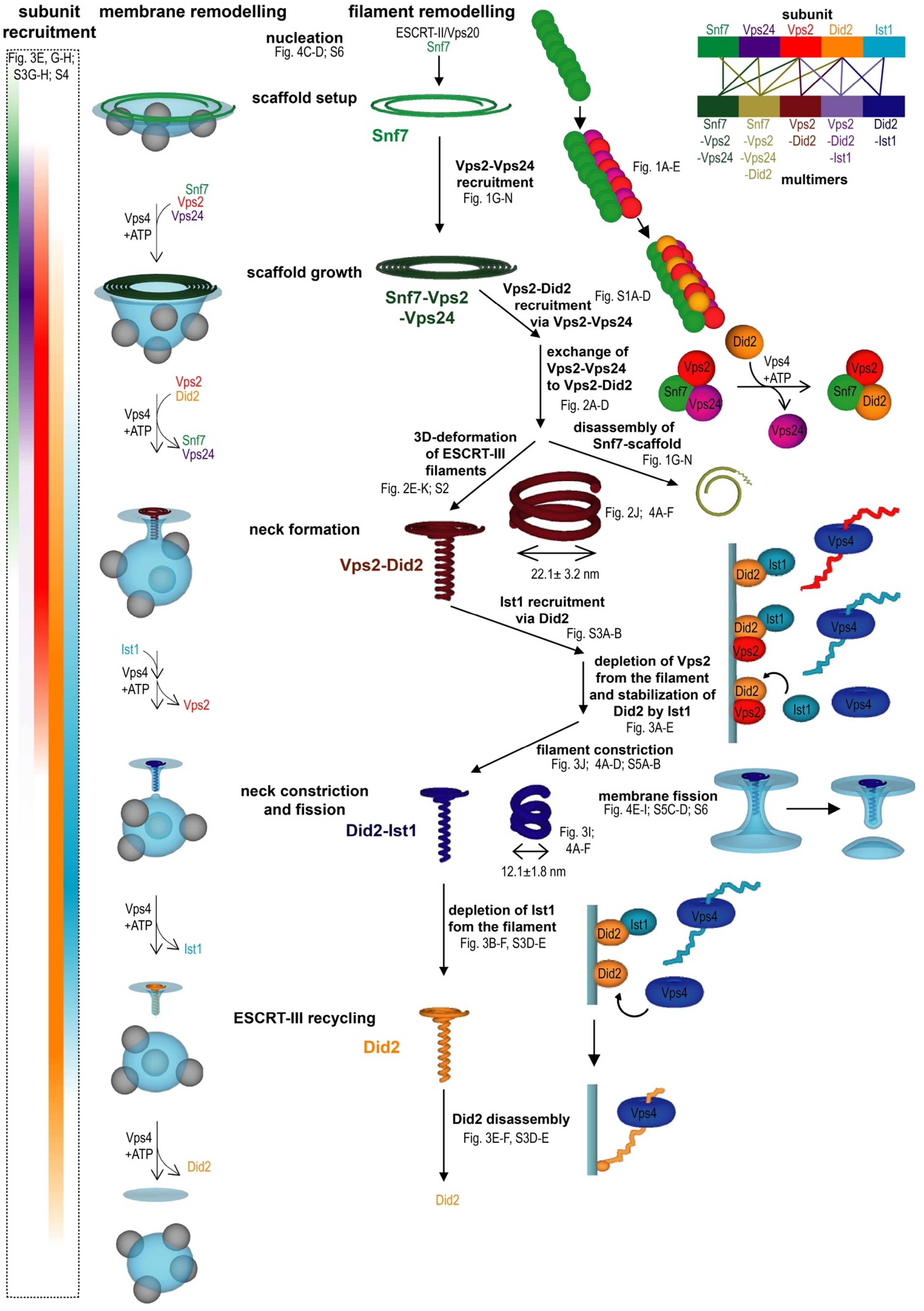

subunit recruitment
membrane remodelling
filament remodelling
Fig. 3E, G-H; S3G-H; S4
nucleation
Fig. 4C-D; S6
ESCRT-II/Vps20
Snf7
scaffold setup
Snf7
Vps2-Vps24 recruitment
Fig. 1G-N
Fig. 1A-E
subunit
Snf7 Vps24 Vps2 Did2 Ist1
Snf7 -Vps2 -Vps24
Snf7 -Vps2 -Vps24 -Did2
Vps2 -Did2
Vps2 -Did2 -Ist1
Did2 -Ist1
multimers
Snf7
Vps2
Vps24
Vps4 +ATP
scaffold growth
Snf7-Vps2 -Vps24
Vps2-Did2 recruitment via Vps2-Vps24
Fig. S1A-D
exchange of Vps2-Vps24 to Vps2-Did2
Fig. 2A-D
Vps2
Did2
Vps4 +ATP
Snf7
Vps24
3D-deformation of ESCRT-III filaments
Fig. 2E-K; S2
disassembly of Snf7-scaffold
Fig. 1G-N
Fig. 2J; 4A-F
22.1± 3.2 nm
neck formation
Vps2-Did2
Ist1 recruitment via Did2
Fig. S3A-B
depletion of Vps2 from the filament and stabilization of Did2 by Ist1
Fig. 3A-E
Ist1
Vps4 +ATP
Vps2
filament constriction
Fig. 3J; 4A-D; S5A-B
neck constriction and fission
Did2-Ist1
Fig. 3I; 4A-F
12.1±1.8 nm
membrane fission
Fig. 4E-I; S5C-D; S6
Vps4 +ATP
Ist1
depletion of Ist1 fom the filament
Fig. 3B-F, S3D-E
ESCRT-III recycling
Did2
Vps4 +ATP
Did2
Did2 disassembly
Fig. 3E-F, S3D-E
Did2

**Fig. S7. Model of the coupling between ESCRT-III recruitment sequence and membrane remodeling activities.**
